# eIF3d and eIF3e mediate selective translational control of hypoxia that can be inhibited by novel small molecules

**DOI:** 10.1101/2025.05.29.656739

**Authors:** Stephen C. Purdy, Kate Matlin, Christopher Alderman, Amber Baldwin, Natasha Shrivastava, Somnath Dutta, Kristofor J. Webb, Arthur Wolin, Dillon P. Boulton, Jyoti Kapali, John D. Landua, Michael T. Lewis, M. Cecilia Caino, James C. Costello, William Old, Xiang Wang, Rui Zhao, Heide L. Ford, Neelanjan Mukherjee

## Abstract

Exposure to hypoxia is linked to increased cellular plasticity and enhanced metastasis; effects which are primarily attributed to the transcriptional activation of large gene programs downstream of hypoxia inducible factors (HIFs). However, translational effects in hypoxia, that likely precede transcriptional effects, have remained largely unexplored. Using ribosome-profiling, we uncovered a selective translational response in acute hypoxia that is eIF3d/eIF3e-dependent and controls downstream hypoxic responses including HIF1a accumulation and cellular invasion. We further demonstrated that eIF3e copy number and an eIF3e-expression signature are associated with worsened outcomes for breast cancer patients. Finally, we identified a class of novel small molecules that target eIF3e specifically, reducing the translational response to hypoxia and to ER stress, another stressor that is dependent on eIF3d/eIF3e-mediated translation. Our data uncover critical functions for eIF3d/eIF3e in the hypoxic response and identify a potential means to inhibit stress-induced translation, and potentially plasticity and metastasis, mediated by eIF3e.

## Introduction

Despite numerous advances in cancer therapies, breast cancer remains the second leading cause of cancer-related deaths among women^1, 2^. Importantly, nearly all breast cancer-related deaths are attributed to metastatic disease^3^, yet a means to target metastasis remains elusive. As cancer cells metastasize, they are exposed to numerous stressors that they must overcome to survive, grow, and metastasize^4^. Recently, the plasticity of mRNA translation has been found to play a critical role in this adaptation^5, 6^. Furthermore, translational reprogramming is a major driver of therapeutic resistance leading to poor prognosis^5, 7^. Translation dysregulation has long been appreciated in cancer initiation and growth. MAPK and mTOR pathways are often highly activated in tumors and largely contribute to tumor cells’ ability to meet the increased protein synthesis demand during oncogenesis^8–10^. Furthermore, alterations in expression of ribosomal proteins and eukaryotic Initiation Factors (eIFs) are common in cancer, with many eIFs having defined oncogenic roles^8^.

Alterations in the translational response to stress can occur independent of transcriptional alterations, resulting in more immediate phenotypic responses. Thus, rapid translational changes are vital for the adaptation to stress^11, 12^. Many stressors are experienced by metastatic cells, such as nutrient deprivation and hypoxia, which cause canonical cap-dependent translation to decrease^13^. A reduction in translation allows cells to conserve energy and building blocks (e.g. amino acids) by limiting the high energy demands of protein synthesis^12^. However, while canonical cap-dependent translation is diminished under various stressors, subsets of mRNAs are selectively translated via non-canonical mechanisms of translation. Selective translation of subsets of mRNAs can be critical for survival of cells in response to stress, and thus may be a major contributor to tumor progression and metastasis. Recently, members of the eIF3 complex have been implicated in selective translation during stress^14–17^.

The eIF3 complex is composed of 13 subunits, providing a large scaffold that is essential for the formation of the 48S preinitiation complex and subsequent translation initiation^18–21^. While the scaffolding function of eIF3 is vital to canonical translation^21^, separable functions have been ascribed to particular subunits of the eIF3 complex that allow for very precise regulation of canonical and non-canonical translation^14–16, 21–32^. For example, eIF3d is now known to bind the 5′ m7G cap of specific mRNAs with stem loop structures in the 5’UTR^23^. This cap binding activity allows eIF3d to promote or repress translation of specific mRNAs^22, 23^. Such noncanonical eIF3d-mediated translation is upregulated during cellular stress, specifically during glucose deprivation and chronic ER stress to promote survival^14, 15^. Additionally, eIF3d is known to promote the epithelial-to-mesenchymal transition (EMT) and may enhance breast cancer metastasis; though direct evidence that eIF3d mediates metastasis is lacking^33^.

The direct binding partner of eIF3d, eIF3e, has also been implicated in cancer, and like eIF3d, regulates translation of a subset of mRNAs in response to various stressors. However, the direct mechanism of eIF3e’s selective translational regulation remains unknown^34^. The role of eIF3e in tumor progression is complex; it has been reported to be either tumor suppressive or promotional, dependent on context^35–39^ ^40–43^. Nonetheless, both eIF3d and eIF3e have described tumor promotional roles, and their ability to regulate selective translation may be critical to tumor progression. While several studies have elucidated functions for eIF3d or eIF3e under basal conditions, few studies have focused on both proteins, particularly during cellular stress, to uncover their overlapping and distinct functions, which would provide important regulatory insights.

One of the most common stressors within the tumor is hypoxia. Hypoxia occurs in 50-60% of solid tumors and is well-known to promote metastasis via multiple mechanisms^4^. A majority of the tumor promotional effects of hypoxia are attributed to the downstream transcriptional regulation by hypoxia-inducible factors (HIFs)^4^. Under normoxic conditions, HIF1a and HIF2a are constitutively expressed, but rapidly degraded by oxygen-dependent prolyl hydroxylases (PHDs). However, during low oxygen conditions, PHDs are inhibited which allow for the rapid accumulation of HIFlZJ proteins. HIFlZJ proteins can then bind to HIF1β (also known as ARNT), and act as pleiotropic transcription factors to regulate the expression of genes involved in angiogenesis, survival, metabolism, and proliferation, amongst other properties^44^. Thus, transcriptional reprogramming in response to hypoxia has been well documented^45, 46^; however, the role of non-canonical translational in response to hypoxia and the factors governing it are largely unexplored.

Herein, we determine the contributions of eIF3d and eIF3e to translational regulation under both normoxic and hypoxic conditions. In normoxia, we identify shared and unique mRNAs that are regulated by eIF3d and eIF3e. We demonstrate that the acute response to hypoxia is predominately regulated translationally, with little changes in RNA levels. We find that the number of mRNAs translationally regulated by eIF3d and eIF3e is dramatically increased in hypoxia compared to normoxia, and that both subunits of the eIF3 complex are required for hypoxic-induced invasion, demonstrating their role in selective translation in a reduced oxygen environment mediates phenotypes associated with metastasis. Furthermore, we identify an acute translational hypoxic response that requires eIF3d and eIF3e. We find that eIF3e copy number gain and RNA-signatures strongly correlate with worsened overall survival in breast cancer. Importantly, we identify a class of novel small molecules that target eIF3e, inhibiting the acute hypoxic response as well as hypoxia-induced invasion. We further show that this novel class of inhibitors targets stress-induced translation of ATF4 in response to ER stress, another stress response mediated by eIF3d/eIF3e. Taken together, we demonstrate a critical role for eIF3d and eIF3e in the hypoxic response, and identify first-in-class inhibitors of eIF3e. Such compounds will not only serve as important research tools to understand the function of eIF3e in stress-induced translation, but also may be starting points for developing novel therapeutic interventions for metastasis.

## Results

### eIF3e and eIF3d have overlapping and distinct functions

To delineate overlapping and distinct functions of eIF3d and eIF3e (Fig. 1A), we performed ribosome-profiling (Ribo-seq) in parallel with RNA-seq in a metastatic breast cancer cell line, MCF7-SIX1^47, 48^, after siRNA-mediated knockdown (KD) of eIF3e or eIF3d. Briefly, Ribo-seq identifies the positions of ribosome on RNAs by isolating and sequencing the ribosome protected fragments (RPFs)^49^ (Fig. 1B). Furthermore, one can calculate translational efficiency (TE) of an individual RNA by comparing the ratio of RPFs to RNA levels and ask how these change across conditions to identify mRNAs subject to translational regulation. Biological replicates for both RNA-seq and Ribo-seq were highly reproducible (RNA-seq R=0.98, Ribo-seq R=0.96) (Fig. S1A). All Ribo-seq data sets exhibited the expected ∼29 nt RPF lengths (Fig. S1B), enrichment in the coding sequence (Fig. S1C), frame preference (Fig. S1D), and three nucleotide periodicity (Fig. S1E). We observed the expected reduction in eIF3e and eIF3d protein levels after their respective depletion (Fig. 1C, Fig. 1A depicts the eIF3 complex to show the relationship of eIF3d and e). While eIF3e KD reduced eIF3d protein levels, similar to what has been observed in other systems^50^, it did not decrease the mRNA levels or RPFs of eIF3d (Fig. S1F). These data suggest that eIF3e does not control eIF3d levels via transcription, mRNA decay, or translation, but that it may instead stabilize eIF3d protein post-translationally. Since eIF3d protein levels decrease with eIF3e KD, loss of eIF3d may be responsible for at least a portion of the mRNA translational changes observed with KD of eIF3e.

**Figure 1:**
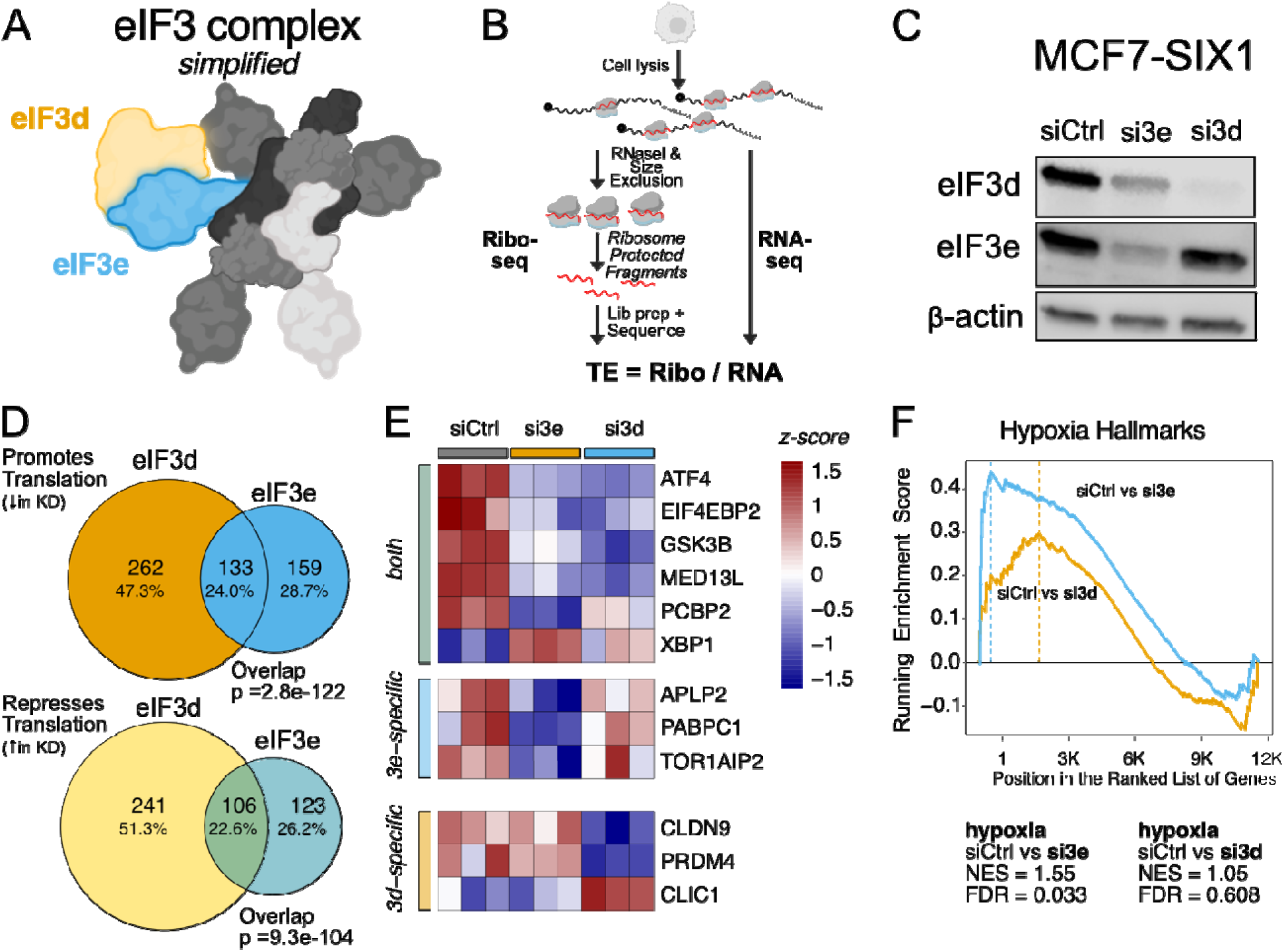
Overlapping and distinct functions of elF3d and elF3e. **A.** Diagram of **a** simplified elF3 complex with only elF3d and 3e labeled for clarity **B.** Outline of the experimental design of Ribo-seq and RNA-seq. **C.** siRNA knockdown of elF3d and elF3e in MCF7-SIX1 cells followed by western blot analysis. **D.** Venn diagram of the overlapping and distinct differentially translated mRNAs after elF3d and elF3e KD. P value calculated using Fisher’s Exact Test. **E.** Heatmap of specific mRNAs that are shared or specific to elF3d vs elF3e. Values are z-scores of TE. **F.** Gene set enrichment analysis for the hypoxia hallmark dataset when elF3e and elF3d are knockdown.

To identify the mRNAs translationally regulated by eIF3d/e, we calculated TE changes upon knockdown (KD) of eIF3d/e compared to the control siRNA^51^. Loss of eIF3d or eIF3e resulted in significantly altered TE of 742 or 521 mRNAs, respectively (Fig. 1D). Similar numbers of mRNAs were upregulated (repressed by eIF3d/e) as downregulated (promoted by eIF3d/e) upon eIF3d/e KD. There was a statistically significant overlap of 239 mRNAs regulated by both eIF3d and eIF3e (133 promoted and 106 repressed) (Fig. 1D). Moreover, the direction and degree of translational changes were concordant for mRNAs regulated by either eIF3e or eIF3d (R^2^=0.21) (Fig. S1G). While some overlap of transcripts regulated by eIF3d or eIF3e would be anticipated as KD of eIF3e decreased eIF3d protein levels, we also found eIF3e and eIF3d to regulate distinct transcripts. We identified transcripts regulated by both eIF3d and eIF3e and transcripts regulated specifically by either eIF3d or eIF3e and visualized representative transcripts (Fig. 1E). We identified similar pathways enriched or depleted for TE changes upon eIF3e and eIF3d depletion (Table S1). Interestingly, mRNAs encoding proteins in the “Hypoxia” hallmark gene set were significantly enriched with eIF3e KD and to a lesser extent with eIF3d KD (Fig. 1F).

### eIF3e and eIF3d are necessary for the acute translational hypoxic response

Given the enrichment of the hypoxia hallmark gene set (Fig. 1F) and that both eIF3d and eIF3e have been associated with various stress responses^14, 15, 25, 52–54^, we decided to further examine their role in the hypoxic response. Because the tumor promotional roles of hypoxia have largely been attributed to the transcriptional response downstream of HIF1a^4, 55^, we first asked if eIF3d or eIF3e influence HIF1a expression in response to hypoxic conditions (4 hours at 1% O_2_). We observed that the induction of HIF1a protein levels in response to hypoxia is significantly attenuated by depletion of either eIF3d or eIF3e in numerous cell lines (Fig. 2A-C). Based on this finding, we tested whether eIF3d and eIF3e are involved in the translational response to hypoxia. Specifically, we performed Ribo-seq in MCF7-SIX1 breast cancer cells +/- eIF3d or eIF3e KD under acute hypoxic (1% O_2_ for 1 hour) vs normoxic conditions (Fig. 2D). We chose an early time point after hypoxia induction (1 hour) to examine the translational changes that occur prior to the many anticipated transcriptional changes downstream of hypoxia-dependent HIF1a induction. RNA-seq and Ribo-seq replicates were again highly correlated in this experiment (R=0.96 and R= 0.99, respectively) (Fig. S2A). We identified almost 30x more statistically significant changes in RPFs (478 mRNAs increased and 807 mRNA decreased) compared to changes in RNA levels (19 mRNAs increased and 26 mRNAs decreased) after one hour of hypoxia (Fig. 2E). More mRNAs exhibited a loss, rather than a gain, of ribosome association, consistent with diminished canonical translation observed in hypoxia^56^ (Fig. 2E). Using these data, we generated an acute translational hypoxic signature, comprising of 358 mRNAs with significant alterations in TE after one hour of hypoxia (Table S1). Importantly, these translational changes were almost completely abolished with eIF3e KD (Fig. 2F) or eIF3d KD (Fig. 2G). Interestingly, while we observed decreases in HIF1a levels with eIF3d and eIF3e KD even at the 1-hour time point (Fig. S2B) the TE for *HIF1*a was not significantly changed, indicating that eIF3d/3e do not directly regulate HIF1a translation in acute hypoxia (Fig. S2C). Taken together, our data indicate that translation is the major early form of gene regulation in response to acute oxygen deprivation and that eIF3e and eIF3d are required for this acute translational hypoxic response, as well as for hypoxia-dependent HIF1a induction.

**Figure 2:**
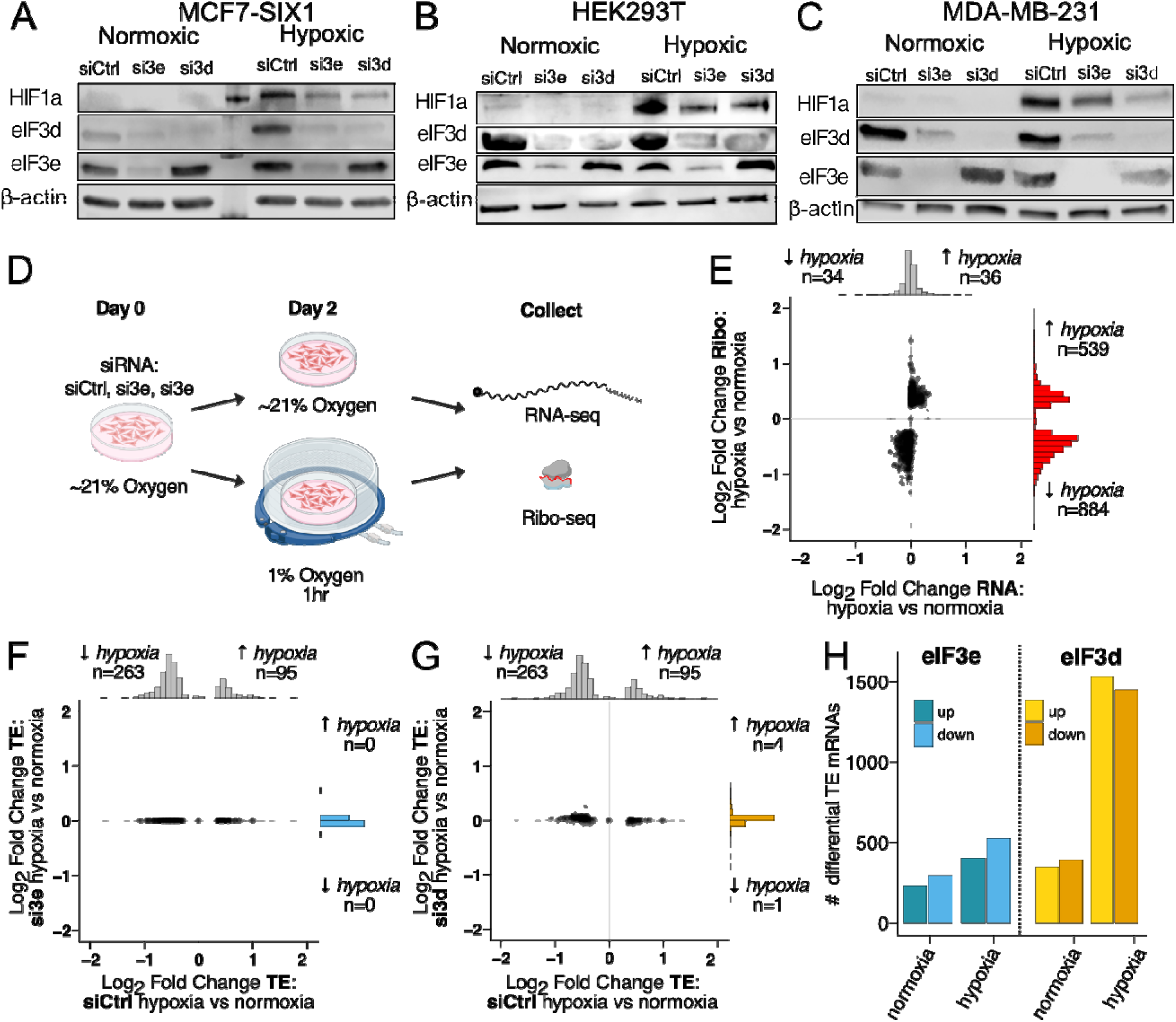
elF3e and elF3d are required for the acute translational hypoxic response. **A-C.** Western blot analysis showing HlFla after elF3d and elF3e knockdown in nonnoxic and hypoxic (4 hours at 1% O2) conditions. **D.** Schematic outlining the experimental design of Ribo-seq/RNA-seq in both normoxic and hypoxic conditions. **E.** Log, fold changes of RPFs (y-axis) and RNA levels (x-axis) comparing siCtrl cells in normoxia vs hypoxia. Only mRNAs with a statistically significant difference (adj p val < 0.05) are shown. **F.** Comparison of TE changes from normoxia to hypoxia in siCtrl cells (x-axis) vs si3e cells (y-axis) **G.** or si3d cells (y-axis). Only mRNAs with a statistically significant TE difference (adj p val < 0.05) are shown. **H.** Barplot of the number of differentially translated mRNAs after elF3e and elF3d KD in normoxia and hypoxia.

We next turned our attention to the full set of mRNAs regulated by eIF3d and eIF3e in both hypoxic and normoxic conditions, rather than solely those mRNAs regulated in the acute translational hypoxic response. In normoxic conditions, eIF3d and eIF3e depletion led to 724 and 521 translationally regulated mRNAs, respectively (Fig. 2H). The number of mRNAs translational regulated by eIF3d and eIF3e depletion in hypoxic conditions were 2991 and 924, respectively. The additional 2249 and 403 mRNAs translationally regulated by eIF3d and eIF3e depletion, respectively, in hypoxic conditions indicates enhanced activity of both eIF3d and eIF3e in response to hypoxia.

### eIF3e and eIF3d regulate induction of hypoxia and hypoxia-induced invasion

Since eIF3e and eIF3d are required for the acute translational hypoxic response, we wanted to explore the functional consequence of their loss to hypoxia-associated phenotypes. To this end, we utilized a previously developed hypoxia fate-mapping system cultured in 3D^57^. In this system, MDA-MB-231 cells were transduced with two plasmids that enable the cells to permanently switch from expressing dsRED during normoxic conditions to expressing GFP during hypoxic conditions^57^. This fluorescent switch is controlled by a Cre recombinase that is transcriptionally regulated via a promoter containing HIF-responsive elements (HREs). Therefore, Cre is only able to cut out the dsRED gene and allow GFP expression when HIF1 is induced in hypoxic conditions^57^. We refer to the hypoxia fate-mapping cells (HFM) as 231HFM. To enable hypoxia to occur in a more natural setting, we embedded the 231 HFM tumorspheres in a mixture of matrigel and collagen. Over time, we observed that the tumorspheres formed a hypoxic core, as evidenced by the induction of GFP (Fig. 3A). As anticipated due to reductions in HIF1a levels (Fig. 2C), both eIF3d and eIF3e KD decreased the ratio of GFP/RFP observed in the tumorspheres over time (Fig. 3A&B). Importantly, we also observed that eIF3d KD and eIF3e KD decreased tumorsphere invasion, when examined over time (Fig. 3C). These data are of interest, as the hypoxic response has been linked to invasion^58^. In line with this finding, our data show that the GFP/dsRed ratio is significantly correlated with increased invasion (Fig. 3D), and that this correlation is lost with both eIF3d and eIF3e KD. These data demonstrate that eIF3d and eIF3e are necessary to mediate the invasive phenotype in response to hypoxia.

**Figure 3:**
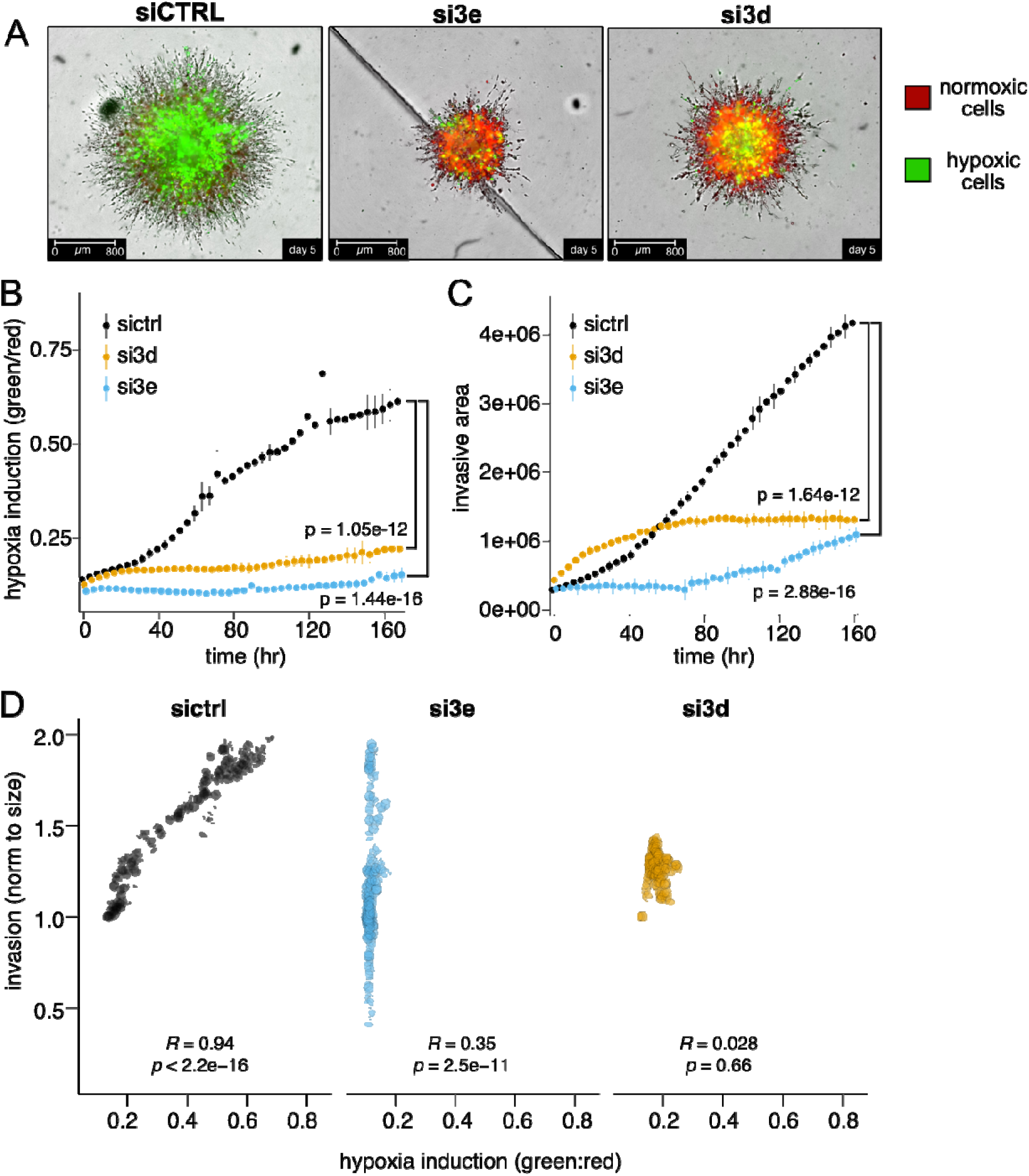
Loss of elF3d and elF3d decreases HIF1a-mediate effects and tumorsphere invasion. **A.** Representative images of 231HFM cells grown in tumorspheres embedded in matrigel and collagen I +/- elF3d and elF3e KD. **B.** Fluorescence ratio of GFP:dsRed over time +/- elF3d and elF3e KD. **C.** Invasive area of the 231 tumorspheres overtime +/- elF3d and elF3e KD. For B and C, statistical significance was calculated using a longitudinal mixed effects model in which si3e and si3d were both compared to sictrl. **D.** Scatterplot of the normalized Invasive area (x-axls) and the GFP:dsRED ratio (y-axls) for sictrl (left), sl3d (middle), and si3e (right). Spearman correlation coefficient and p-value denoted for each.

### The gene signature associated with eIF3e, but not eIF3d, predicts poor prognosis in breast cancer patients

Hypoxia is known to promote tumor progression and correlate with worse prognosis among breast cancer patients^4, 55, 59^. Since our data show that eIF3d and eIF3e promote the hypoxic response in breast cancer cells, we asked whethe*r EIF3D* or *EIF3E* were dysregulated in large breast cancer patient cohorts, such as METABRIC (1980 patients), in a manner correlated with survival. We found that 3% and 46% of patients exhibited gain/amplification in the *EIF3D* or *EIF3E* alleles (Fig. S3A), respectively. Interestingly, *EIF3E* gain/amplification significantly correlated with worse prognosis, while *EIF3D* gain/amplification did not (Fig. 4A-B). However, since the chromosomal location of *EIF3E* (8q23) is very close to that of *cMYC* (8q24), which is amplified in ∼70% of human malignancies and acts as a prominent oncogene^60^, the prognostic value of EIF3E may be driven by *cMYC* co-amplification. Therefore, we developed *EIF3D* and *EIF3E* signatures from RNA-seq in MCF7SIX1 cells with eIF3d or eIF3e KD in hypoxia (p < 0.05 and LFC > 1 or <-1), to identify a prognostic signature specific to *EIF3D* and *EIF3E* independent of *cMYC*. We stratified patients based on the enrichment of *EIF3D* and *EIF3E* RNA signatures. Enrichment of the *EIF3E* signature was associated with worsened survival compared to patients depleted of the *EIF3E* signature (Fig. 4C). The eIF3d signature did not correlate with overall survival (Fig. S3B). Next, we asked how a previously published 42-gene mRNA hypoxia signature conserved across breast cancer cell lines^61^ stratified patient survival. As expected, enrichment of the hypoxia signature was associated with worsened patient survival in the METABRIC dataset^62–64^ (Fig. 4D).

**Figure 4:**
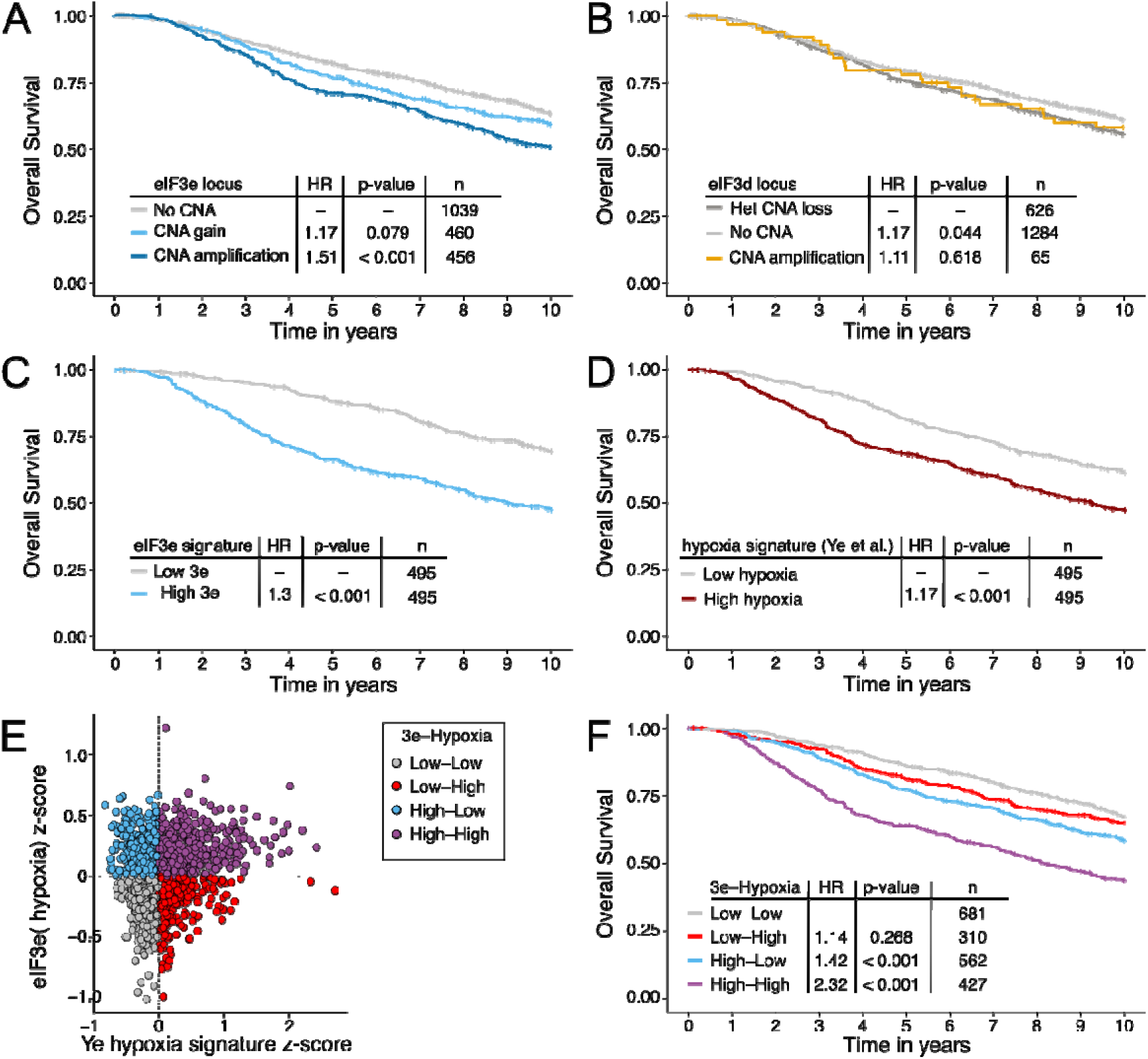
elF3e-associated RNA signature predicts poor prognosis. **A&B.** Overall survival rates of METABRIC patients (n=1980) with breast tumor **(A)** elF3e gains/amplifications or **(B)** elF3d gains/amplifications compared to no copy number alterations (CNA) in the METABRIC dataset. **C.** Overall survival of patients in the METABRIC dataset stratified by our elF3e RNA-seq signature. For clarity only the first and fourth quartile are shown. **D.** Overall survival of patients from the METABRIC dataset stratified by a 42-gene hypoxic signature. **E.** Categorizing patients based on elF3e RNA-seq signature z-score and their hypoxia z-score. **F.** Overall survival rates of patients in the METABRIC dataset stratified by the combination of elF3e and hypoxia signature enrichment/depletion. The p-values and hazard ratios were calculated using a Cox proportional hazards regression where each group was compared to the control group (e.g. no CNA for elF3e, het CNA loss for elF3d, bottom 25% for signatures).

Since enrichment of the conserved hypoxic signature and the eIF3e signature were both associated with worsened outcomes (Fig. 4B-C) and the acute hypoxic translational response was dependent on eIF3e (Fig. 2F), we investigated potential dependencies between hypoxia and eIF3e signatures for patient survival. To this end, we grouped patients into four categories based on high/low hypoxia and high/low eIF3e signature enrichment (Fig. 4E). This analysis demonstrates that the high eIF3e/high hypoxia group correlates with the worst survival outcomes, whereas the low 3e/low hypoxia group correlates with the best survival outcomes (Fig. 4F). While not statistically significant, the high eIF3e/low hypoxia group trended with worsened survival when compared to the low eIF3e/high hypoxia group (Fig. 4F), suggesting that eIF3e may have additional, hypoxia-independent roles in breast cancer that mediate outcomes. Together, these data provide compelling evidence for the relevance of targeting eIF3e in breast cancer.

### Small molecules targeting eIF3e

We previously demonstrated that a small molecule, NCGC00378430 (abbreviated **8430**), reduces metastasis in an MCF7-SIX1 xenograft model^48^. While 8430 decreased the interaction between SIX1 and EYA2 (two known tumor promotional molecules), we also observed that levels of both proteins were decreased, and thus the direct target of this molecule remained unclear^48^. To identify the direct target of 8430, we performed isothermal shift assays (ITSAs) on cells treated with 8430 or a vehicle control (DMSO) and found that eIF3e was the most significantly stabilized protein in the presence of 8430 (Fig. 5A). To independently confirm stabilization of eIF3e with 8430, we performed cellular thermal shift assays (CETSAs) on MCF-7SIX1 lysates and found that 8430 stabilized eIF3e at 53 °C, while eIF3d was not stabilized (Fig. 5B). After this discovery, we synthesized closely related compounds to 8430, and found that an analog compound (209), also stabilizes eIF3e in CETSAs (Fig. 5B). We then performed isothermal dose-response fingerprint (ITDRF) experiments, in which a dose-response curve is generated to calculate half-maximal effective concentrations (EC_50_). The calculated EC_50_s are 13.6 µM and 18.3 µM for 8430 and 209, respectively (Fig. 5C-F). These data suggest that 8430 and 209 bind to eIF3e and provide a potential explanation for the prevention of metastasis by 8430^48^, via inhibition of eIF3e.

**Figure 5:**
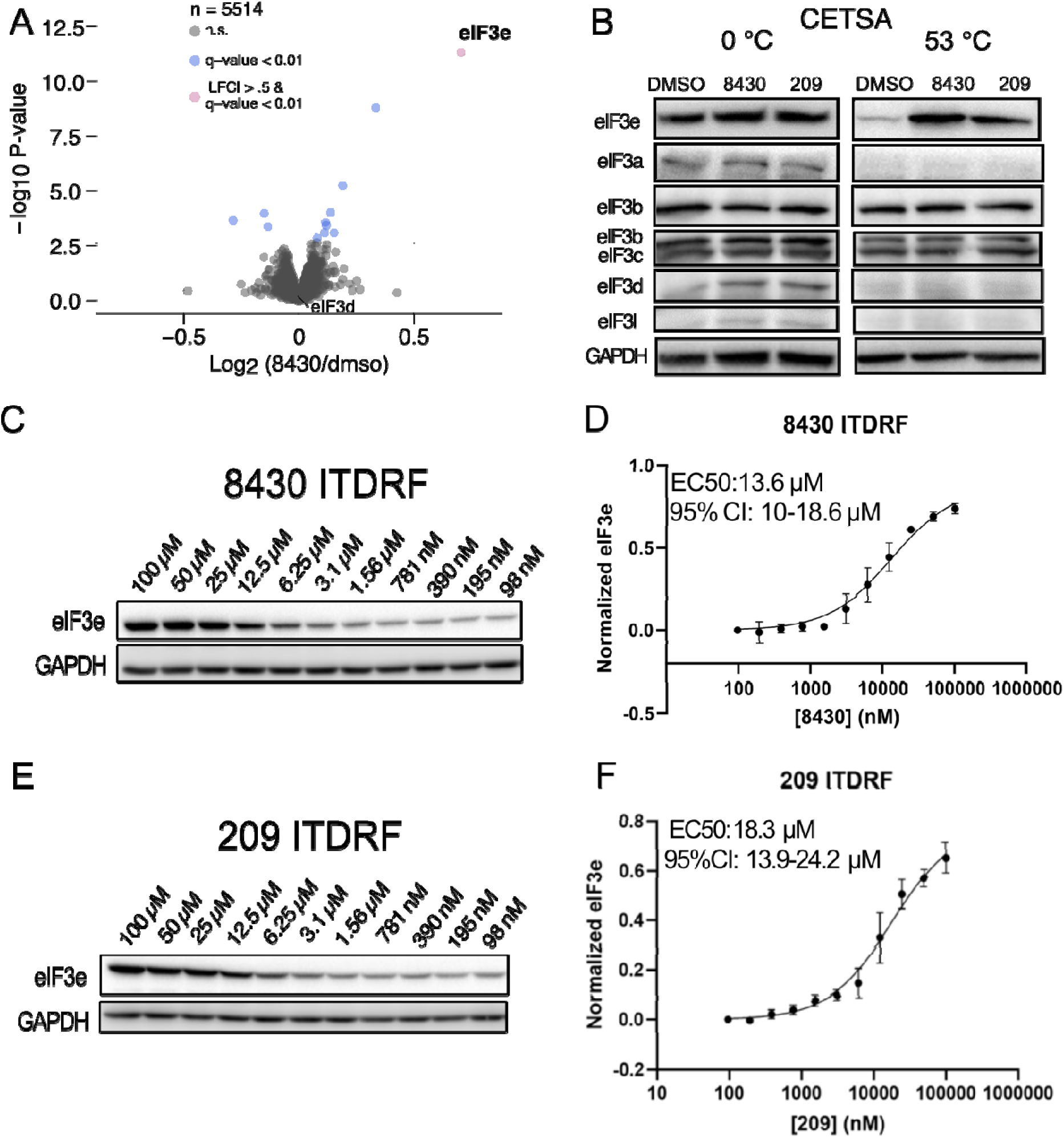
Compounds 8430 and 209 specifically stabilize elF3e protein in thermal shift assays. **A.** Volcano plot of the Isothermal shift assay (ITSA) Iog2 fold change (x-axis) and -log 10 p value (y-axis) **B.** Cellular thermal shift assay (CETSA) followed by western blot showing protein levels of elF3e and multiple other elF3 subunits in addition to a loading control or GARDH. **C&D.** Dose response of 8430 for an Isothermal dose-response fingerprint (ITDRF) assay. **E&F.** Dose response of 209 for an Isothermal dose-response fingerprint (ITDRF) assay.

### Pharmacologic inhibition of eIF3e inhibits the acute translational hypoxic response

To determine whether our novel class of eIF3e targeting small molecules inhibit eIF3e-associated hypoxic responses, we first looked at HIF1a levels after decreasing O_2_ levels to 1% for 4 hours, +/- 8430 and 209 treatment. As observed with both eIF3d and eIF3e KD, treatment of both MCF7-SIX1 and HEK293T cells with 8430 and 209 led to reduced HIF1α induction in response to hypoxia (Fig. 6A-B, S6A). Given the promising similarities, we performed Ribo-seq in normoxic or hypoxic conditions for 1 hour +/- treatment with DMSO (vehicle) or compound 209, which was chosen due to the larger reduction of HIF1a levels when compared to 8430 (Fig. 6A-B). RNA-seq and Ribo-seq replicates were again highly correlated in normoxia (R=0.99 and R=0.93, respectively) and hypoxia (R=0.99 and R=0.95, respectively (Figure S4A-B). As observed previously in our siCTRL cells (Fig. 2E), acute hypoxia caused many changes in ribosomal footprints, with minimal changes to transcript levels, indicating translation as the major regulatory response at this timepoint (Figure 6C). In line with our eIF3e KD results (Fig. 2F), the acute hypoxic translational response was largely ablated in MCF7-SIX1 breast cancer cells treated with 209 (Fig. 6D). Similarly, we identified more mRNAs with significantly changed TEs after 209 treatment in hypoxic conditions when compared to normoxic conditions (Fig. 6E). Neither the RNA levels nor RPFs for HIF1a changed in response to 209, again suggesting that eIF3e does not directly regulate the translation of HIF1a (Fig. S4C). Interestingly, no significant overlap was observed between mRNAs translationally regulated by 209 treatment and those regulated by eIF3e or eIF3d in normoxic conditions (Fig. 6F). In contrast, we observed a highly significant overlap between mRNAs regulated by eIF3e and 209 in hypoxic conditions (Fig. 6G, blue), whereas eIF3d and 209 showed a more moderate, yet significant, overlap with specifically the downregulated mRNAs in hypoxic conditions (Fig. 6G, yellow). These results further suggest that our compound is on-target and inhibits eIF3e function. Taken together, our data demonstrate that there is a unique, eIF3d/e translational mechanism in hypoxia that can be specifically inhibited pharmacologically with a small molecule targeting eIF3e.

**Figure 6:**
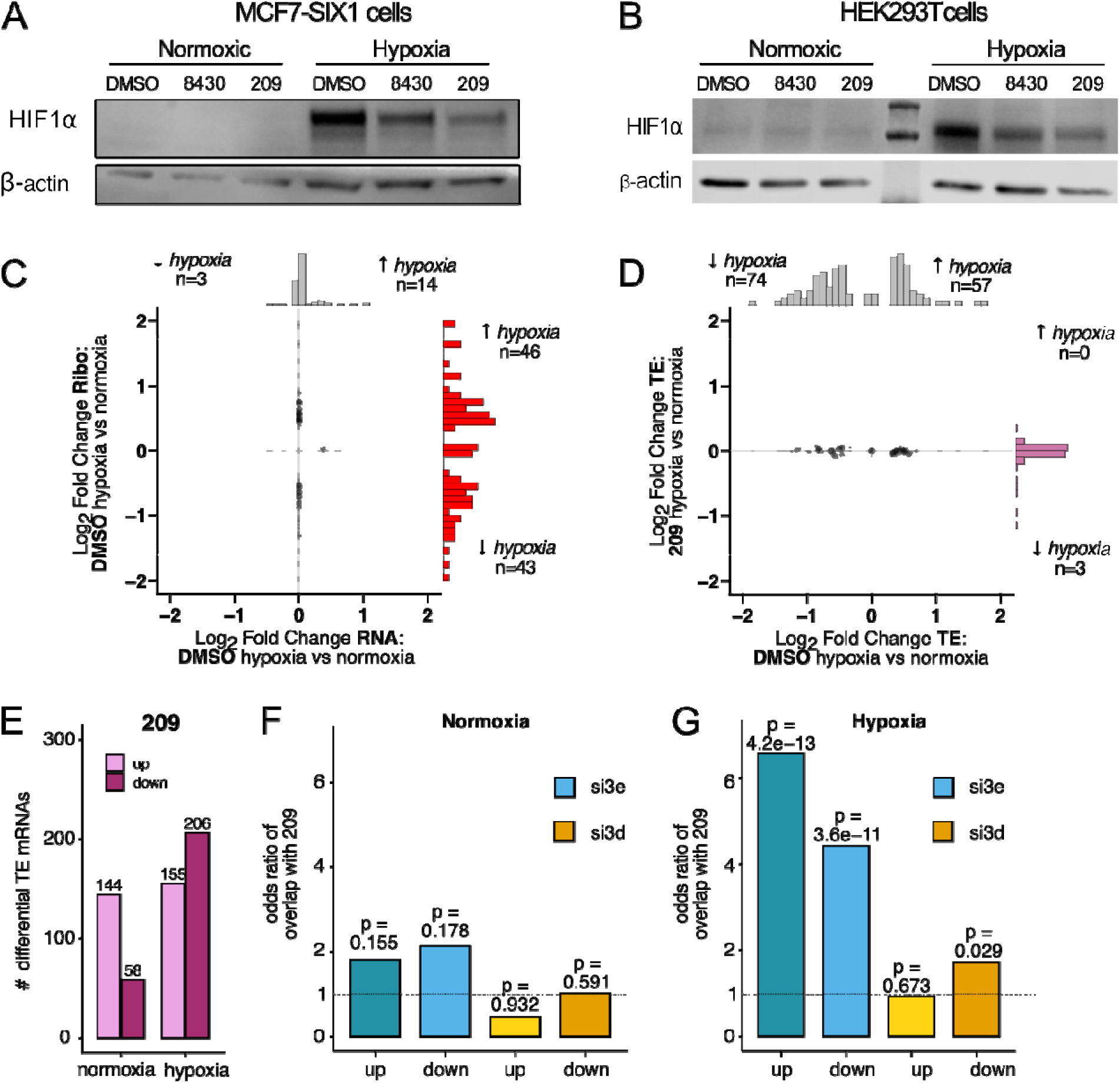
Small molecule compounds, 8430 and 209, inhibit the acute translational hypoxic response. **A&B.** Western blot analysis showing HIF1a after 8430 and 209 treatment (24 hour pre-treatment) in normoxic and hypoxic (4 hours at 1%O2) conditions. C. Log_2_ fold change of RPFs (y-axis) and RNA levels (x-axis) comparing DMSO treatment in normoxia vs hypoxia. Each dot represents a mRNA transcript. **D.** Comparison of significant TE changes from normoxia to hypoxia in DMSO treated cells (x-axis) to TE changes from normoxia to hypoxia after treatment with a209 (y-axis). **E.** Barplot of the number of differentially translated mRNAs with 209 treatment in normoxia and hypoxia. Barplot of the odds ratio of the overlap of mRNA with TE changes from a209 treatment compared to those with TE changes from elF3e or elF3d KD In **F.** normoxia and G. hypoxia. P value calculated using Fisher’s Exact Test.

### Pharmacologic inhibition of eIF3e prevents hypoxic-related phenotypes

Because our novel eIF3e-targeting compounds inhibited HIF1a induction and selective translation in response to acute hypoxia, we again utilized the 231 HFM tumorsphere model to determine whether hypoxia induced phenotypes could be pharmacologically inhibited. Since the tumorsphere assays take place over a week time period, we first assessed the stability of both 8430 and 209 over time. Our data demonstrate that compound 209 degrades over time in our culture media, whereas 8430 is stable over the 7 days used in our tumorsphere assays (Fig. S5). Thus, we used 8430 to examine whether pharmacological targeting of eIF3e could inhibit hypoxia-induced phenotypes in 3D over time. Importantly, 8430 significantly decreased the GFP:dsRED ratio within these tumorspheres over time (Fig. 7A-B), similar to eIF3e KD. Furthermore, 8430 decreased tumorsphere invasion over the time course of this assay (Fig. 7C). Again, the GFP:dsRED ratio was correlated with invasion (R value=0.96); however, this correlation is decreased with the addition of 8430 (R value=0.47) (Fig. 7D). These data demonstrate the ability of our eIF3e targeting compound (8430) to functionally inhibit hypoxia-induced invasion, and provides further evidence that 8430 is acting on-target given the similarities to eIF3e KD.

**Figure 7:**
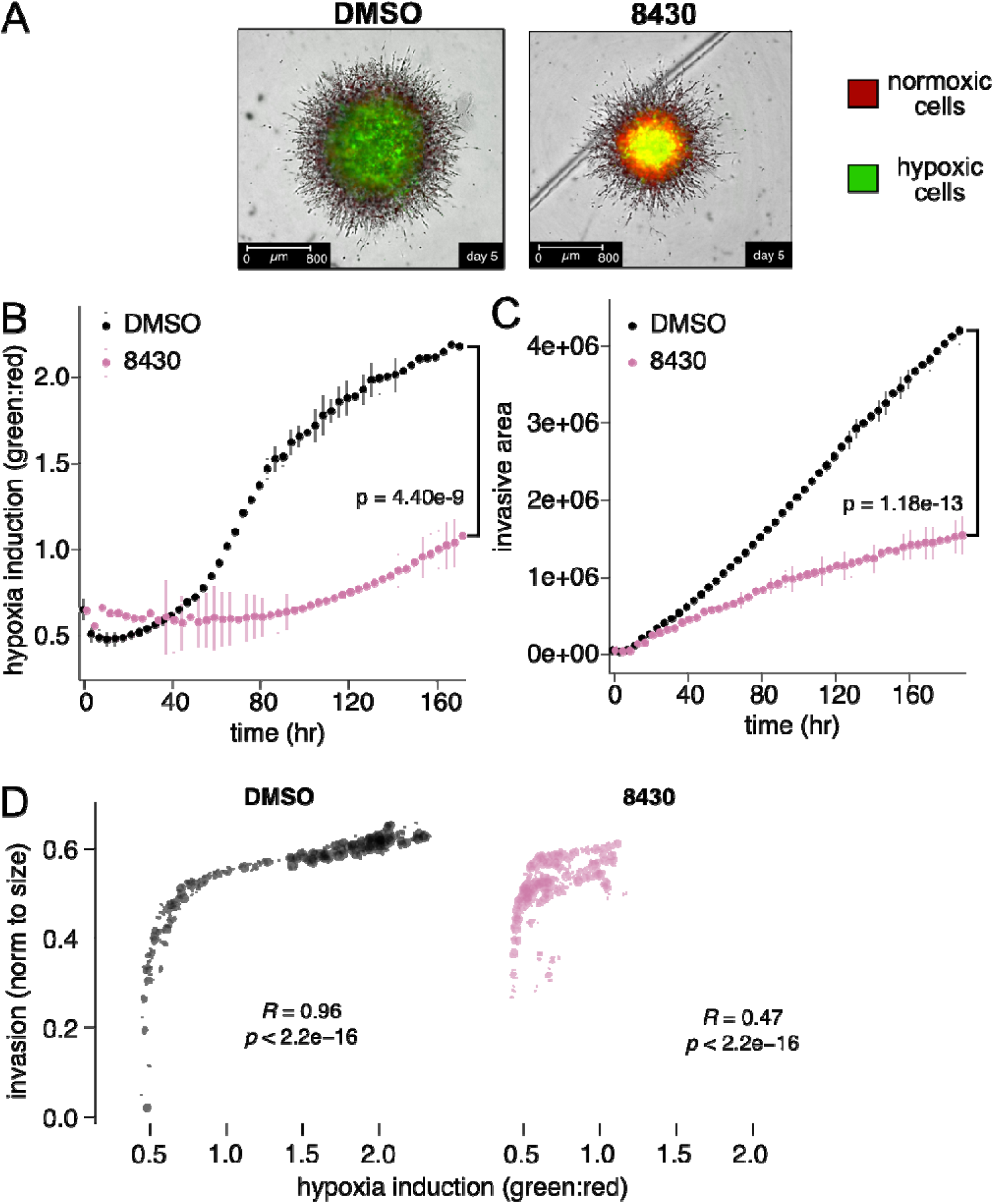
8430 inhibits HI Fla-mediate effects and tumorsphere invasion. **A.** Representative images of 231HFM cells grown in tumorspheres embedded in matrigel and collagen I +/- 8430 treatment. **B.** Fluorescence ratio of GFP:dsRed over time +/- 8430 treatment. **C.** Invasive area of the 231 tumorspheres over time *+1-* 8430 treatment.For B and C, statistical significance was calculated using a longitudinal mixed effects model comparing 209 to DMSO. **D.** Scatter plot comparing the normalized invasive area (y-axis) and the GFP: dsRED ratio (x-axis) in DMSO (left) and 8430 (right) treatment. Spearman correlation coefficient and p-value denoted for each.

### Response to additional cellular stressors requires eIF3e and eIF3d, and can be inhibited with our novel eIF3e-targeting compounds

Because we demonstrate a role for both eIF3e and eIF3d in the hypoxic response, and because both molecules have been implicated in response to a variety of stressors^14, 15, 25, 52–54^, we further explored whether eIF3d and eIF3e are cooperating to regulate additional stress induced responses, and whether our eIF3e targeting small molecule could inhibit these responses. Of interest, we observed that eIF3e *and* eIF3d both promoted the TE of the *ATF4 mRNA,* a known master transcriptional regulator of the integrated stress response (ISR)^65, 66^, in our original experiment under normoxic conditions (Fig. 1E). To further explore the role of eIF3e and eIF3d in regulating ATF4 protein expression, we treated a variety of cell lines with thapsigargin (TG) or tunicamycin (TM)—two potent inducers of ISR^17^—to induce ATF4 translation. We verified that induction of ATF4 is abrogated with both eIF3e KD and eIF3d KD in multiple cell lines (Fig. S6A-B). Of note, we noticed abnormally high levels of ATF4 basally in our MCF7-SIX1 cells. We found these cells were uniquely sensitive to siRNA transfection which activated the ISR (Fig. S6C). This finding likely explains why we found ATF4 as an mRNA translationally regulated by eIF3e and eIF3d KD without addition of an ER stressor. Other cell lines that we tested not sensitive to siRNA transfection, also retained loss of ATF4 in both eIF3e and eIF3d KD cell lines after TG and TM treatment. Importantly, both 8430 and 209 also significantly attenuated the induction of ATF4 expression after TG and TM treatment, similar to eIF3d KD and eIF3e KD (Fig. S6D-E). Taken together, these data demonstrate that 8430 and 209 can inhibit the other stress-responsive phenotypes that are dependent upon eIF3e.

## Discussion

In this study, we demonstrate that eIF3d and eIF3e are required for cancer cells to respond to hypoxia through regulating a selective translational program, and that this translational program is critical for induction of HIF1a and the phenotypic response to hypoxia. We further demonstrate that that eIF3e RNA signatures are associated with poor overall survival in breast cancer patients. Importantly, we have developed first-in-class compounds that target eIF3e and inhibit the acute translational hypoxic response, resulting in reduced invasion in response to hypoxia. As we have previously demonstrated that one of these novel compounds inhibits metastasis in vivo^48^, our results suggest that our first-in-class inhibitors may reduce metastasis at least in part via altering the response to hypoxia through eIF3e.

Hypoxia is a well-known stressor that can promote tumor progression and metastasis^67^. During stress, cells often downregulate mTOR signaling, thus diminishing canonical eIF4F-mediated translation. However, a subset of mRNAs is still translated via non-canonical translation programs, which is thought to be vital to cellular adaptation. However, the hypoxia-induced mechanism by which non-canonical translation is mediated remains underexplored. Herein, we find that eIF3d and eIF3e are critical for the initial translational response to hypoxia. While eIF3d and eIF3e regulate translation in normoxic conditions, there are 1.8x and 4.8x more translationally regulated mRNAs by eIF3e and eIF3d, respectively, in hypoxia, suggesting an enhancement of this translational program during stressed conditions. This translational response precedes the well-defined pleiotropic transcriptional response of HIF1a—the response that is attributed to the tumor promotional effects of hypoxia. Furthermore, hypoxia-induced HIF1a accumulation is abrogated by the loss of eIF3d and eIF3e, suggesting that inhibition of the hypoxic translational response will also attenuate further downstream HIF1a-mediated transcriptional responses. Interestingly, we did not observe changes in *HIF1a* mRNA or RPFs with loss of eIF3d or eIF3e, suggesting that the change in HIF1a downstream of eIF3d and eIF3e is a function of altered protein stability. Furthermore, we did not see significant changes in HIF1a regulators such as VHL—the main regulator of HIF1 stability^68^. Although our study did not uncover a translational effect of eIF3d or eIF3e on HIF1a, a previous report demonstrated that eIF3e loss results in decreased *HIF1*a translation in glioblastoma cells^16, 54^. However, this study utilized a hypoxia mimetic, CoCl_2_, to stabilize HIFa proteins, rather than low oxygen levels, which may account for the differences in regulation of HIF1a observed when compared to our study. Furthermore, it is possible that HIF1a is translationally regulated at a later timepoint (>1 hour).

This work expands on previous studies that have found a selective eIF3d-dependent translational response that occurs when various stressors (glucose deprivation, integrated stress response activators) inhibit canonical translation, allowing translation of a subset of mRNAs—such as cJUN and ATF4—to promote survival^14–16, 69^. Furthermore, eIF3e has been shown to promote resistance to radiotherapy, and is overexpressed in patients with recurrent glioblastoma tumors post radiotherapy, suggesting that eIF3e is required for adapting to this stress^54^. These data add to the mounting evidence for the crucial role of these translation factors for adaptation to a wide variety of stressors, including microenvironmental stressors and current therapies.

We found that our eIF3e-associated RNA signature, as well as a previously described hypoxia RNA signature^70^ both predict poor prognosis in the METABRIC breast cancer dataset. When taken together with our observation that eIF3e KD inhibits pro-metastatic phenotypes such as invasion, these findings underscore the potential value of developing small molecules to target eIF3e as a novel therapeutic strategy. In our current study, we demonstrate that novel small molecule compounds, 8430 and 209, stabilize eIF3e but not other eIF3 components, suggesting that they specifically bind eIF3e. These compounds recapitulate phenotypes seen in eIF3e KD cells, such as preventing the acute translational hypoxic response, HIF1a induction, HIF1-mediated dsRED:GFP switch, and invasion in 3D, further suggesting these compounds are on-target. Importantly, we have previously shown that 8430 can inhibit metastasis in our in vivo MCF7-SIX1 xenograft model^48^. In that work, we demonstrated that 8430 reduced expression of a pro-metastatic transcription factor, SIX1, and its cofactor, EYA2, respectively. 8430 was able to reverse SIX1-mediated transcriptional changes, SIX1- and TGFb-induced epithelial to mesenchymal transition (EMT), and SIX1-induced metastasis^48^. However, we did not observe a decrease in the TE of SIX1 or EYA2 with eIF3e KD or with 209 treatment, suggesting that eIF3e indirectly regulates these pro-metastatic transcription factors. Nevertheless, it is exciting that 8430 targets eIF3e and inhibits metastasis in vivo. To conclusively demonstrate that our eIF3e targeting compounds are reducing metastasis through eIF3e specifically, it will be important to perform genetic experiments in which eIF3e is depleted and animals are treated in vivo with 8430. Also, future biophysical studies are needed to identify the precise mechanism of action of our compounds on eIF3e, and to further optimize the compounds to improve drug-like characteristics such as potency and pharmacokinetic parameters. Nonetheless, our compounds can serve as tools to understand the function of eIF3e in various cellular contexts.

Tumor cells encounter countless stressors that are constantly fluctuating as they proliferate, deplete resources, and metastasize throughout the body^71^. During these times of cellular stress, tumor cells are in a state of vulnerability and must adapt to survive. The ability of tumor cells to adapt engenders a plasticity that is challenging to target therapeutically^5, 6, 72^. Therefore, identifying commonalities between mechanisms of adaptation to various stressors and exploiting these vulnerabilities could enable the development of novel therapeutic strategies for targeting cancer cells at multiple stages of tumor progression, including metastasis^73^. Devising novel means to inhibit selective translation may be one such approach. Indeed, many studies have shown the importance of mRNA translation for tumor plasticity and progression^5, 72, 74^. To this end, various translation inhibitors have been developed but have limited use in the clinic often due to toxicity or lack of efficacy^75, 76^. However, the development of strategies to specifically inhibit stress-induced translation—which is generally highly active in malignant tissues— could uncover key vulnerabilities to limit tumor plasticity with minimal toxicities. Our studies and others provide a growing body of literature that suggest eIF3d and eIF3e are critical for adaptation to a variety of different stressors^14, 15, 25, 52–54^, indicating these proteins are viable therapeutic targets for this approach.

## Methods

### Cell lines and cell culture

Cells were cultured in incubators at 37 with 5% CO_2_ and ∼21% O_2_ (atmospheric). MCF7SIX1 cells (Clone A13) were generated as previously described^47^. The MDA-MB-231 cells were a generous gift from Dr. Daniele Gilkes^57^. HEK293T cells were acquired from the University of Colorado Anschutz Medical Campus Cell Technology shared resource. The MCF7SIX1 and HEK293T cells were cultured Dulbecco’s High Glucose Modified Eagle Medium (DMEM, Hyclone SH30022.01), with which 10% fetal bovine serum (FBS, Corning 35-010-CV), and 2 mM L-glutamine (L-glut, Hyclone SH30034.02) were supplemented. The MDA-MB-231 cells were cultured with Dulbecco’s High Glucose Modified Eagle Medium (DMEM, Hyclone SH30022.01), with which 5% fetal bovine serum (FBS, Corning 35-010-CV), 2 mM L-glutamine (L-glut, Hyclone SH30034.02), were supplemented. All cell lines were STR profiled at the end of the study to ensure identity, and were also tested for mycoplasma every 6 months with the MycoAlert Mycoplasma Detection Kit (Lonza LT07–118). Cell lines were passaged after thawing prior to use in experiments and were re-thawed periodically. 8430 and 209 were resuspended in DMSO to a working solution concentration of 20 mM. All experiments with 8430 and 209 were used at 20 mM (1:1000 dilution) with DMSO as a vehicle control. TG and TM were used at 100 nM and 2.5 mg/mL, respectively, with DMSO as a vehicle control.

### siRNA transfections

ON-TARGET*plus* siRNA SMARTPools were ordered specifically for eIF3e (cat #: L-010518-00-0010) and eIF3d (cat #: L-017556-00-0010) from Horizon Discovery. ON-TARGET*plus* non-targeting pool (cat # D-001810-10) was always used as a control. siRNA transfections occurred in 25% OPTIMEM Reduced serum media and 75% full growth media (cat # 31985062). siRNAs were added to the media mixture at a concentration of 7.5 nM for 5 minutes prior to the addition of 1.875 mL Lipofectamine RNAI MAX transfection reagent (cat # 13778150) per 1 mL of media. siRNA:lipofectamine mixtures were incubated for 20 minutes before adding it directly onto the cell culture overnight. siRNA knockdowns were assessed 48-72 hours after the addition of the siRNAs.

### Western Blotting

Cells were washed with PBS then either frozen in the plate (dry), or lysed fresh with RIPA buffer. Cells were detached and whole-cell protein extracts were isolated by lysing cell pellets using RIPA buffer. Collected protein lysates (30–60 μg) were separated using polyacrylamide gel electrophoresis, and size-separated proteins were then transferred to methanol-activated PVDF membranes followed by the blocking step using 5% non-fat milk in TBST (20 mM Tris HCl, pH 7.5, 150 mM NaCl, 0.1% Tween-20). Next, membranes were incubated with indicated primary antibodies (1:1000) overnight at 4°C. The following day, membranes were washed with TBST and secondary antibodies (1:5000) were applied in 5% non-fat milk in TBST. Finally, membranes were washed and examined using chemiluminescence followed by imaging using the Odyssey Fc Imaging System (LI-COR Biosciences) or the Azure 600 Gel imaging system (Azure biosystems).

### RNA-seq and Ribosome profiling

Ribosome profiling was performed as described previously with a few modifications^77^. The lysis buffer contained 100 ug/mL cycloheximide. Following lysis and clarification by centrifugation, total RNA was quantified using the Qubit RNA BR assay kit. A portion of the lysate was reserved as input sample for RNASeq preparation. Total RNA was extracted using the DirectZol RNA Microprep kit (ZYMO Research) following the manufacturer’s instructions with on-column DNase I treatment. Ribosomal RNA was depleted from total RNA using our previously described RNaseH-based method^78^, then input into the KAPA RNA Hyper prep RNAseq library prep kit (Roche) according to manufacturer’s instructions with dual-index adapters, followed by quality check of the final libraries by Qubit dsDNA HS assay and Tapestation High Sensitivity D1000 screentape. Libraries were pooled and submitted for paired-end 150 bp sequencing on the Illumina NovaSeq6000 platform to a depth of 40 million paired end reads per library.

The remaining lysate was used as input into the ribosome footprinting reaction with 750 U RNase I (Invitrogen Ambion). The digestion was stopped with SUPERaseIN followed by recovery of monosomes with equilibrated Microspin S-400 HR (GE) columns. Trizol LS was added to the monosome-containing elution and RNA isolated with the ZYMO Directzol microprep kit with on-column DNase I treatment. The isolated RNA was depleted of rRNA using the siTOOLs riboPOOLs kit for ribosome profiling. The samples were resolved on a 15% TBE-Urea PAGE gel and ribosome protected fragments (RPFs) 28-29 nt in size were extracted and recovered. RPFs were treated with T4-PNK then used as input into the Qiagen QIAseq miRNA library prep kit with provided single index adapters. Libraries were pooled and submitted for paired-end 150 bp sequencing on the Illumina NovaSeq 6000 S4 platform to a depth of 50 million paired end reads per library (Novogene Corporation Inc.). Adapters were trimmed with Cutadapt^79^. UMIs were trimmed and collapsed using UMI-tools^80^. Salmon^81^ was used for quantifying transcript levels from RNA-seq and Ribo-seq libraries using GENCODEv26 transcriptome assembly. Briefly, Salmon data were imported using the tximport^82^. Transcript expression was quantified using Salmon v1.6 (12). Quality of ribosome sequencing data, including ribosome footprint length, p-site positioning, and three-nucleotide periodicity, was assessed using Ribowaltz^83^. All RNA-seq and Ribo-seq reads were compared using Pearson correlation, by which an outlier was identified and removed from downstream analysis (Hypoxia, si3e, Rep B, RNA-seq). Differential expression and translation was calculated using the deltaTE approach^51^. The odds ratio and p-value for the overlap between genes was performed using a Fisher’s exact test. The pipeline and all downstream analysis, which was performed in R with custom scripts, are available at https://github.com/mukherjeelab/2024_eIF3e_hypoxia/.

### Hypoxic conditions

To provide a hypoxic environment, cells were placed in either a modular hypoxic chamber and filled with a premixed gas comprised of 1% O_2_, or the cells were placed in a BioSpherix hypoxic chamber flushed with N_2_ and CO_2_ gas. The O2 and CO2 concentrations in the BioSpherix chamber were maintained at 1% and 5%, respectively, using a carbon dioxide & oxygen controller (BioSpherix). These conditions were maintained constant throughout the course of the experiments. After hypoxic incubation was completed, quick lysis (<3 min) of the cells was necessary to prevent reoxygenation from affecting our results.

### Event-Free survival analysis and eIF3e/eIF3d/hypoxia signatures

eIF3e and eIF3d RNA signatures were made by comparing the LFC of RNA in siCtrl cells compared to KD cells in hypoxia. Genes included in each signature had a log_2_ fold change <-1 or >1 and an adjusted p value <0.05. mRNA microarray data for 2509 primary breast tumor samples were downloaded from cBioPortal for Cancer Genomics^84, 85^ specifically from the METABRIC Breast Cancer study^62, 63^. Duplicate microarray data for patient entries were merged by taking the mean of the z-score for each gene. Patients with missing microarray data were removed. After preprocessing, we performed downstream analysis on the remaining 1980 patients.

For copy number variation analysis, patients were grouped by eIF3e or eIF3d copy number. For signature analysis, patients were assigned to groups based on the top quartile of enrichment (top 25% n=495) and bottom quartile of enrichment i.e. depletion (bottom 25% n=495) for a given signature. To determine statistical differences between groups, each group was compared to the lowest expression group (e.g. no CNA for eIF3e, het CNA loss for eIF3d, bottom 25% for signatures,). Statistics were calculated using a Cox proportional hazards regression analysis.

### Tumorsphere assays

8,000 231HFM cells were plated in each well of a 96-well ULA round bottom plate (Corning cat#7007), centrifuged for 3 minutes at 150 G, then incubated overnight. For the siRNA KD experiments, these were plated 24 hours after siRNA transfection. For 8430 experiments, 8430 (or DMSO) was added to the tumorspheres at time of plating. Matrigel (Corning cat#354234) (10mg/mL) was thawed and kept on ice. Matrigel was mixed 1:1 with full growth media (addition of DMSO or 8430 was added, if applicable). Then, rat tail Collagen I (Corning cat#354249) was added to either 10% total volume (for KD experiments) or 3% of the total volume (8430 experiments) and mixed thoroughly. This mixture of matrigel, media, collagen was carefully pipetted down the side of the wells without disturbing the tumorsphere, then incubated at 37 degrees C for 45 minutes to solidify. Additionally full growth media (with 8430 or DMSO, if applicable), was added on top of the solidified matrigel pad. The plate was then added to the incucyte SX5 (sartorius) for images to be taken at 4x over time. The green and orange optical module were used to measure the fluorescence of the GFP and dsRED, respectively. The brightfield optical module was used for total spheroid size and invasion area.

### Synthesis of 8430 and 209

Detailed description in supplemental material.

### Isothermal Shift Assays

Full description provided in supplemental material. Cell lysates were prepared by suspending frozen cell pellets in 1 mL of lysis buffer: phosphate-buffered saline supplemented with 0.4% Igepal CA-630, a 1X cOmplete™ Mini EDTA-free Protease Inhibitor Cocktail (Sigma #11836170001), and 1X Pierce™ Phosphatase Inhibitor (Thermo Fisher Scientific #A32957), adjusted to pH 7.4. Cells were lysed through sonication. The lysate was then centrifuged at 21,000 g for 15 minutes at 4°C, and the supernatant was collected. Protein concentration was quantified using a bicinchoninic acid (BCA) assay (Thermo Fisher Scientific, Waltham, MA), and the lysate was standardized to a concentration of 5 mg/mL. Cell lysates were treated with compound 8430 at a final concentration of 20 μM, with DMSO adjusted to a 1% final concentration. A vehicle control containing only 1% DMSO was simultaneously prepared.

Isothermal proteome profiling using was performed using the ITSA method, as previously described^86^. A 40 μL aliquot of lysate was dispensed into individually sealable PCR wells (4titude Random Access plate, PN 4ti-0960/RA). The sample plate was equilibrated to room temperature for 2 minutes, sealed and positioned in the thermal cycler for a 3-minute incubation at a constant temperature of 53°C. The plate was then immediately cooled to 4°C, followed by centrifugation at 500 g for 2 minutes to remove condensation.

Protein aggregates were pelleted by transferring PCR tubes to 1.5 mL microcentrifuge tubes and centrifuging at 21,000 g for 30 minutes at 4°C. Supernatants were carefully transferred to fresh low-retention tubes. Protein samples were denatured and cysteines alkylated by adding 50 μL of denaturing buffer (8 M guanidine HCl, 100 mM HEPES (pH 8.5), 10 mM TCEP, and 40 mM 2-chloroacetamide) and heating samples to 90 °C for 10 minutes. After cooling to ambient temperature, 400 μL of cold acetone was added and samples incubated overnight at −20°C.

Samples were centrifuged at 21,000 g for 30 minutes at −10°C. Acetone wash steps were performed twice using 80% cold acetone, with intermediate sonication in a Bioruptor at 4°C. Acetone pellets were dried and resuspended in a digestion master mix: 10% trifluoroethanol, 50 mM HEPES (pH 8.5), Lys-C (2 μg, Wako #129-02541), and trypsin (2 μg, Sigma #T6567). Samples were suspended by sociation and digested for 16 hours at 37°C with constant agitation (1,200 rpm) using an Eppendorf Thermal Mixer C. Peptide labeling utilized the 10-plex TMT reagents (Thermo Fisher Scientific #90110) according to manufacturer protocols.

Peptides were fractionated using an Agilent 1100 HPLC. Mass spectrometric analysis utilized a Waters M-class Acquity system coupled with an Orbitrap Fusion (Thermo Scientific). Peptides were separated using a gradient elution from 3% to 85% acetonitrile over 121 minutes. MS2 spectra were collected using data-dependent acquisition with specified resolution and AGC parameters. Raw mass spectrometry files were processed using MaxQuant (version 1.6.3.323) with the Andromeda search engine. Searches were conducted against a UniProt human protein database, implementing stringent peptide identification criteria. Quantitative analysis employed R (version 3.5.2) with the limma package. Data preprocessing included filtering, quantile normalization, and statistical analysis using linear modeling and empirical Bayes methods. Multiple testing correction was performed using the Benjamini-Hochberg procedure to control false discovery rate.

### Cellular Thermal Shift Assay (CETSA) and Isothermal Dose-Response Fingerprint (IDRF) CETSA

MCF7SIX1 cells were grown to 95% confluency and trypsinized for 5 minutes at 37 °C, neutralized with FBS-containing media and pelleted at 1,000 RPM for 10 minutes at 4 °C. Cell pellets containing 1.2×10^6^ cells were incubated on ice for 30 minutes with 100 µL of lysis buffer containing 50 mM pH 7.5 Tris, 100 mM NaCl, 1 mM DTT, and protease inhibitor cocktail (Thermo Scientific). The cell pellets were then lysed by pulling them through a 23G syringe needle 8 times. Lysed cells were centrifuged at 15,000 RPM for 30 minutes at 4 °C and the supernatant collected. Lysate was treated with either 20 μM of compound in DMSO or 2% DMSO (CETSA) or a serial dilution of compound concentrations (ITDRF CETSA) for 10 minutes at room temperature. The cell lysate was heated at 53 °C for 3 minutes in a thermocycler or incubated at 0 °C on ice and rotated briefly on a microtube rotator to bring down the condensation on the side of the tube. The samples were then transferred to low-adhesion 1.5 mL tubes and centrifuged at 15,000 RPM for 1 hour at 4 °C. 30 µL of supernatant was collected from each tube. Western blots were performed on the resulting supernatants using the following antibodies: α-GAPDH (mouse, GeneTex GT239), α-eIF3a (rabbit, Bethyl A302-002A), α-eIF3b (rabbit, Bethyl A301-760A), α-eIF3c (rabbit, Bethyl A300-377A), α-eIF3d (rabbit, Bethyl A301-758A), α-eIF3e (rabbit, Bethyl A302-984A), α-eIF3k (rabbit, Bethyl A301-762A), and α-eIF3l (rabbit, Bethyl A304-754A).

### Comparative studies of stability between 8430 and 209

#### Thin layer chromatography (TLC)

TLC was carried out on silica plate with 10% ethyl acetate-hexane solvent system. 8430 and 209 were treated with buffer for 1, 5, and 7 days and checked for decomposition after extraction with sodium bicarbonate dichloromethane.

#### Quantitative NMR (qNMR)

qNMR was used to investigate how much of pure 8430 and 209 were present in the buffer solution. Ethelyne carbonate (13 mg for 209, 11 mg for 8430) was considered as internal standard, *d* 4.49 in DMSO-D6 and 4.54 in CDCl3. For 8430 qNMR analysis, *d* 7.20 was considered to be the analyte chemical shift with 2.03 integration, whereas *d* 3.11 was considered with 4.00 integration for 8430 qNMR analysis.

## Acknowledgments

We thank Dexiang Gao for guidance with statistical analyses and Dr. Daniele Gilkes for the hypoxia fatemapping system (231HFM cells).

## Funding

National Institutes of Health grant R35GM147025 (NM)

National Institutes of Health grant R35GM145289 (RZ)

National Institutes of Health grant R01 CA224867 (HLF and MTL)

National Institutes of Health grant CA221282 (HLF and RZ)

National Institutes of Health NRSA T32CA174648 (CA)

National Institutes of Health 5T32CA190216-08 (SCP)

National Institutes of Health T32-GM136444 (KM)

The Dan L. Duncan Comprehensive Cancer Center BCM, P30CA125123 (MTL)

The Cancer Prevention and Research Initiative of Texas BCM, RP170691 (MTL)

Cancer League of Colorado grant AWD-242679-NM (NM and HLF)

This work used the Cell Technologies (RRID:SCR_021982) and Biostatistics and Bioinformatics Shared Resource (RRID:SCR_021983) supported by P30CA046934

## Author contributions

Conceptualization: SCP, KM, RZ, NM, HF

Investigation: SCP, KM, CA, AB, NS, SD, KJW, AW, DPB, JK, JDL

Supervision: MCC, JCC, MTL, WO, XW, RZ, NM, HF

Writing—original draft: SCP, KM

Writing—review & editing: SCP, KM, RZ, NM, HF

## Disclosure and competing interests statement

A patent application has been filed (XW, WO, RZ, HLF) on “Small molecules targeting eIF3e to inhibit tumor growth progression, and metastasis”. Application #:18/839,131. All other authors declare they have no competing interests.

## Data and materials availability

All processed data and code are available at https://github.com/mukherjeelab/2024_eIF3e_hypoxia/. Raw data will be made accessible upon acceptance at GEO and PRIDE, respectively.

## Supplementary Methods

### Synthesis of 209

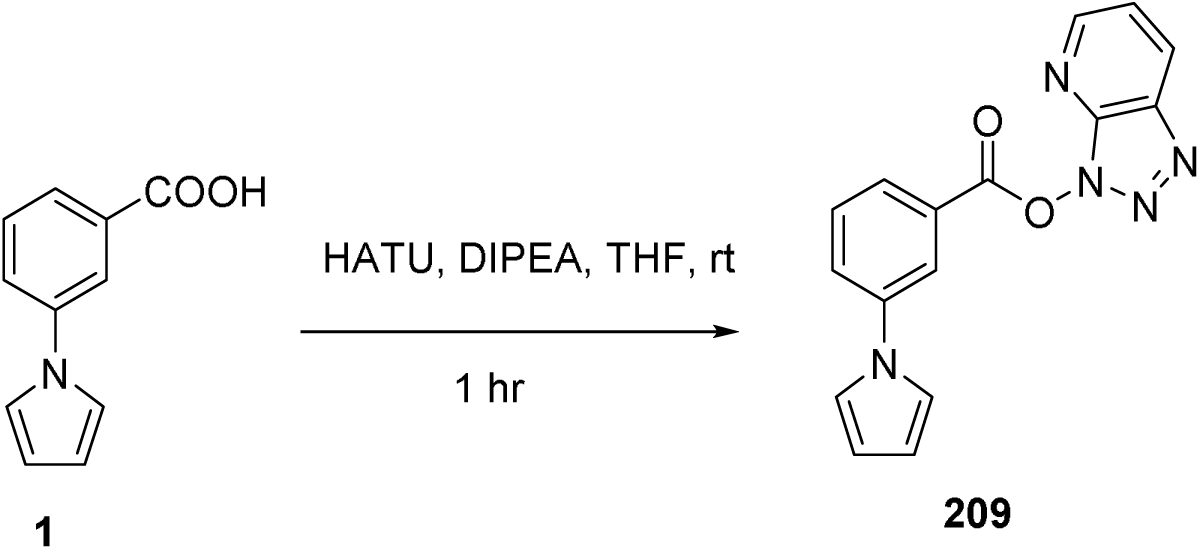

To dry THF (16 mL, 0.2 M) solution of 1 (500 mg, 3.205 mmol) in a flame dried round bottomed flask under argon were added HATU (3.65 g, 9.615 mM), DIPEA (1.67 mL, 9.615 mM) at room temperature. The reaction was stirred for 1 hr. After completion, the reaction mixture was quenched with a saturated NaHCO3 solution (30 mL) and extracted with EtOAc (5x 10 mL). The combined organic layer was dried over anhydrous Na2SO4 and concentrated under reduced pressure. The crude residue was purified by a flash column chromatography on silica using 10% EtOAc-Hexane to afford the product 209 which was titurated with diethyl ether to get the pure while solid (Rf = 0.7, 782 mg, 85% yield).

M.P. 129oC −133oC. 1H NMR (300 MHz, CDCl3) δ 8.77 (dd, J = 4.5, 1.4 Hz, 1H), 8.51 (dd, J = 8.4, 1.4 Hz, 1H), 8.32 (t, J = 1.9 Hz, 1H), 8.20 (dt, J = 7.7, 1.4 Hz, 1H), 7.82 (ddd, J = 8.2, 2.4, 1.1 Hz, 1H), 7.70 (t, J = 7.9 Hz, 1H), 7.51 (dd, J = 8.4, 4.5 Hz, 1H), 7.20 (t, J = 2.2 Hz, 2H), 6.44 (t, J = 2.2 Hz, 2H).

13C NMR (75 MHz, CDCl3) δ 162.10, 151.92, 141.44, 140.70, 135.12, 130.56, 129.70, 127.59, 126.86, 126.13, 122.10, 121.04, 119.20, 111.60.

IR (cm-1): 3150, 3050, 1800, 1600, 1550, 1500, 1360, 1260, 1150, 1050, 990, 790.

HRMS (ESI+) m/z [M+Li]+calcd. for C16H11N5O2: 312.1073; found 312.1048.

### Synthesis of 8430

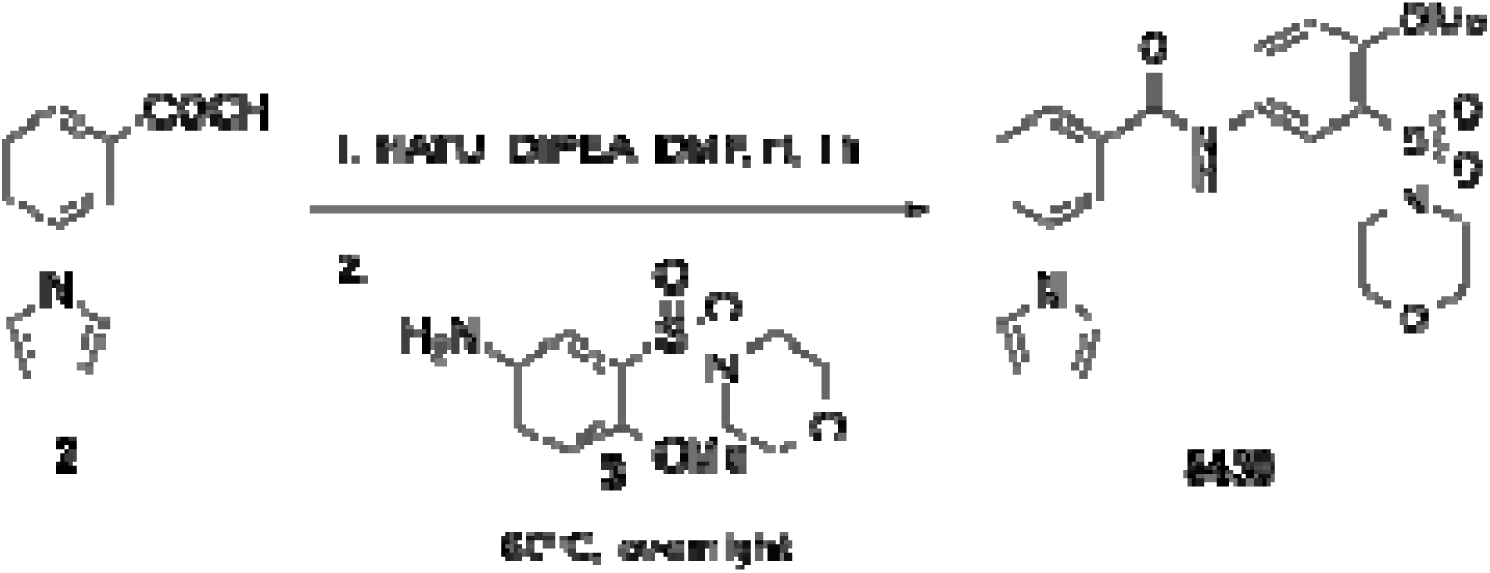

To dry DMF (1.4 mL, 0.2 M) solution of 2 (50 mg, 0.267 mM) in a flame dried round bottomed flask under argon were added HATU (319.36 mg, 0.864 mmol), DIPEA (0.14 mL, 0.864 mM) at room temperature. After 1 hr. 31 was added dropwise and the reaction was heated to 60oC for overnight. After completion, the reaction was quenched with saturated NaHCO3 (10 mL) and extracted with EtOAc (5 mL x 2). The combined organic layer was washed with 1M HCl (10 mL), 1 M NaOH (10 mL), brine (10 mL). The final combined organic layer was dried over anhydrous Na2SO4 and concentrated under reduced pressure. The crude was purified by column chromatography on silica using 50% EtOAc-Hexane as eluent to afford the product 8430 as white solid (Rf = 0.3, 72 mg, 61% yield).

M.P. 210-215oC. 1H NMR (300 MHz, CDCl3) δ 10.47 (s, 1H), 8.23 (d, J = 2.7 Hz, 1H), 8.15 – 8.04 (m, 2H), 7.87 – 7.75 (m, 2H), 7.61 (t, J = 7.9 Hz, 1H), 7.48 (t, J = 2.2 Hz, 2H), 7.31 (d, J = 9.0 Hz, 1H), 6.32 (t, J = 2.2 Hz, 2H), 3.90 (s, 3H), 3.61 (t, J = 4.7 Hz, 4H), 3.10 (dd, J = 5.9, 3.5 Hz, 4H).

13C NMR (75 MHz, CDCl3) δ 169.91, 158.11, 145.14, 141.20, 137.00, 135.12, 132.04, 129.88 (d, J = 21.2 Hz), 128.15, 127.62, 124.42, 123.52, 118.77, 116.02, 71.14, 61.48, 51.05.

IR (cm-1): 3400, 3000, 2950, 2900, 1650, 1580, 1500, 1400, 1350, 1300, 1150, 990, 790.

HRMS (ESI-) m/z [M-H]-calcd. for C21H21N3O5S: 440.5020; found 440.5023.

#### Quantitative NMR (qNMR)

qNMR was used to investigate how much of **8430** and **209** were purely present in the buffer solution. Ethelyne carbonate (13 mg for **209**, 11 mg for **8430**) was considered as internal standard, *δ* 4.49 in DMSO-D6 and 4.54 in CDCl3. For **8430** qNMR analysis, *δ* 7.20 was considered to be the analyte chemical shift with 2.03 integration, whereas *δ* 3.11 was considered with 4.00 integration for **8430** qNMR analysis. Using the formula here (https://www.sigmaaldrich.com/deepweb/assets/sigmaaldrich/marketing/global/documents/101/854/qnmr-brochure-rjo.pdf?srsltid=AfmBOoqpM4uvUVF6NAk5HxeRxmC2vGIbfLxDQ6LXk9Uodbw2mNjbG-LK), **8430** (12.0 mg) showed 93.20% (only the selected protons were showed) of stability continuously from 1 to 5d to 7d with identical ^1^H NMR. **209** (12.0 mg) could not tolerate the buffer and decomposed to show its stability from 96% to 17% (only the selected protons were showed) from 1d to 5d to 7d with identical ^1^H NMR for 5d and 7d. This result concluded that 8430 is more stable than **209** and can intact its structural integrity in that buffer.

### Isothermal Shift Assays

Preparation of lysate—Cell lysates were prepared by suspending frozen cell pellets in 1 mL of lysis buffer: phosphate-buffered saline (137 mM NaCl, 2.7 mM KCl, 10 mM NaHPO4, and 1.8 mM KH2PO4) supplemented with 0.4% Igepal CA-630, a 1X cOmplete™ Mini EDTA-free Protease Inhibitor Cocktail (Sigma #11836170001), and 1X Pierce™ Phosphatase Inhibitor (Thermo Fisher Scientific #A32957), adjusted to pH 7.4. Cell lysis was achieved through mechanical disruption. Briefly, the cell pellet was first suspended by vortexing, then subjected to sonication using a Bioruptor water bath sonicator (Diagenode, Lorne, Australia). The sonication protocol consisted of three 30-second cycles with 30-second intervals between each cycle, performed at 4°C. Following sonication, the lysate was centrifuged at 21,000 g for 15 minutes at 4°C, and the supernatant was collected. Protein concentration was quantified using a bicinchoninic acid (BCA) assay (Thermo Fisher Scientific, Waltham, MA), and the lysate was standardized to a concentration of 5 mg/mL. Prepared samples were immediately utilized for thermal profiling experiments.

Thermal Shift Sample Preparation—Cell lysates were prepared with compound 8430 at a final concentration of 20 μM, with DMSO adjusted to a 1% final concentration. A vehicle control containing only 1% DMSO was simultaneously prepared.

Isothermal proteome profiling using was performed using the ITSA method, as previously described^86^. A 40 μL aliquot of lysate was dispensed into individually sealable PCR wells (4titude Random Access plate, PN 4ti-0960/RA). The experimental workflow involved carefully equilibrating the sample plate to room temperature for 2 minutes before thermal treatment. The plate was then sealed and positioned in the thermal cycler with a heated lid (95°C) for a 3-minute incubation at a constant temperature of 53°C. Immediately following the 3 minute thermal treatment, the plate was immediately cooled to 4°C, followed by centrifugation at 500 g for 2 minutes to remove condensation.

Sample Preparation and Tandem Mass Tag Labeling—Samples were centrifuged at 21,000 g for 30 minutes at −10°C. Acetone wash steps were performed twice using 80% cold acetone, with intermediate sonication in a Bioruptor at 4°C (three cycles, 30-s on, 30 s off). Acetone pellets were dried and resuspended in a digestion master mix: 10% trifluoroethanol, 50 mM HEPES (pH 8.5), Lys-C (2 μg, Wako #129-02541), and trypsin (2 μg, Sigma #T6567). Sonication-assisted resuspension was performed until a homogeneous suspension was achieved. Samples were digested for 16 hours at 37°C with constant agitation (1,200 rpm) using an Eppendorf Thermal Mixer C. Peptide labeling utilized the 10-plex TMT reagents (Thermo Fisher Scientific #90110) according to manufacturer protocols. Briefly, TMT labels were added to individual samples and incubated at room temperature for 1 hour. Labeling was quenched with hydroxylamine, and samples were combined, acidified, and desalted using an Oasis HLB column.

Peptide Fractionation and Mass Spectrometry Analysis—Peptide fractionation employed high-pH reversed-phase chromatography using an Agilent 1100 HPLC. A gradient elution from 5% to 100% acetonitrile was performed over 120 minutes, generating 24 concatenated fractions. Mass spectrometric analysis utilized a Waters M-class Acquity system coupled with an Orbitrap Fusion (Thermo Scientific). Peptides were separated using a gradient elution from 3% to 85% acetonitrile over 121 minutes. MS2 spectra were collected using data-dependent acquisition with specified resolution and AGC parameters.

Computational Data Analysis—Raw mass spectrometry files were processed using MaxQuant (version 1.6.3.323) with the Andromeda search engine. Searches were conducted against a UniProt human protein database, implementing stringent peptide identification criteria. Quantitative analysis employed R (version 3.5.2) with the limma package. Data preprocessing included filtering, quantile normalization, and statistical analysis using linear modeling and empirical Bayes methods. Multiple testing correction was performed using the Benjamini-Hochberg procedure to control false discovery rate.

---

**Figure S1:**
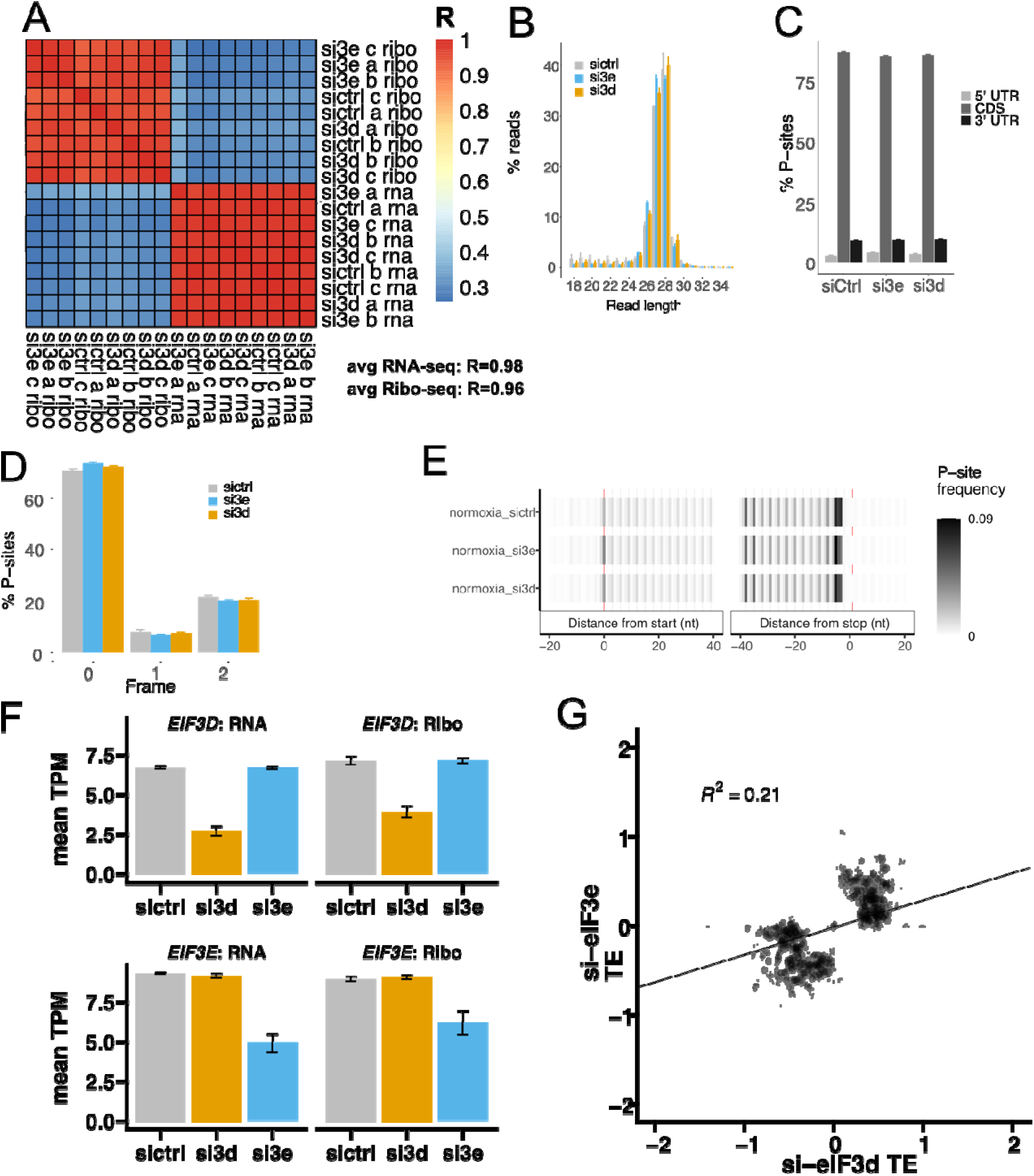
A. Heatmap of Pearson correlation coefficient for normoxic RNA-seq and Ribo-seq Iog2 transcript per million (TPM). B-E. ribosome protected fragments (RPFs) quality control. B-D show the mean and standard deviation of all replicates; similar QC was observed but not shown for all ribo-seq data in manuscript but Is available In Git Repo. B. Read length distribution of RPFs. C. Percent of P-sites annotated to mRNA regions. D. Percent of P-sites in different frames. E. Meta-analysis of P-site positiians relative to start and stop codongs. F. RNA and RPF TPMs of elF3d across samples (top) and elF3e across samples (bottom). G. Log2 fold change of TE values for all protein coding genes after elF3d KD (x-axis) and elF3e KD (y-axis).

**Figure S2:**
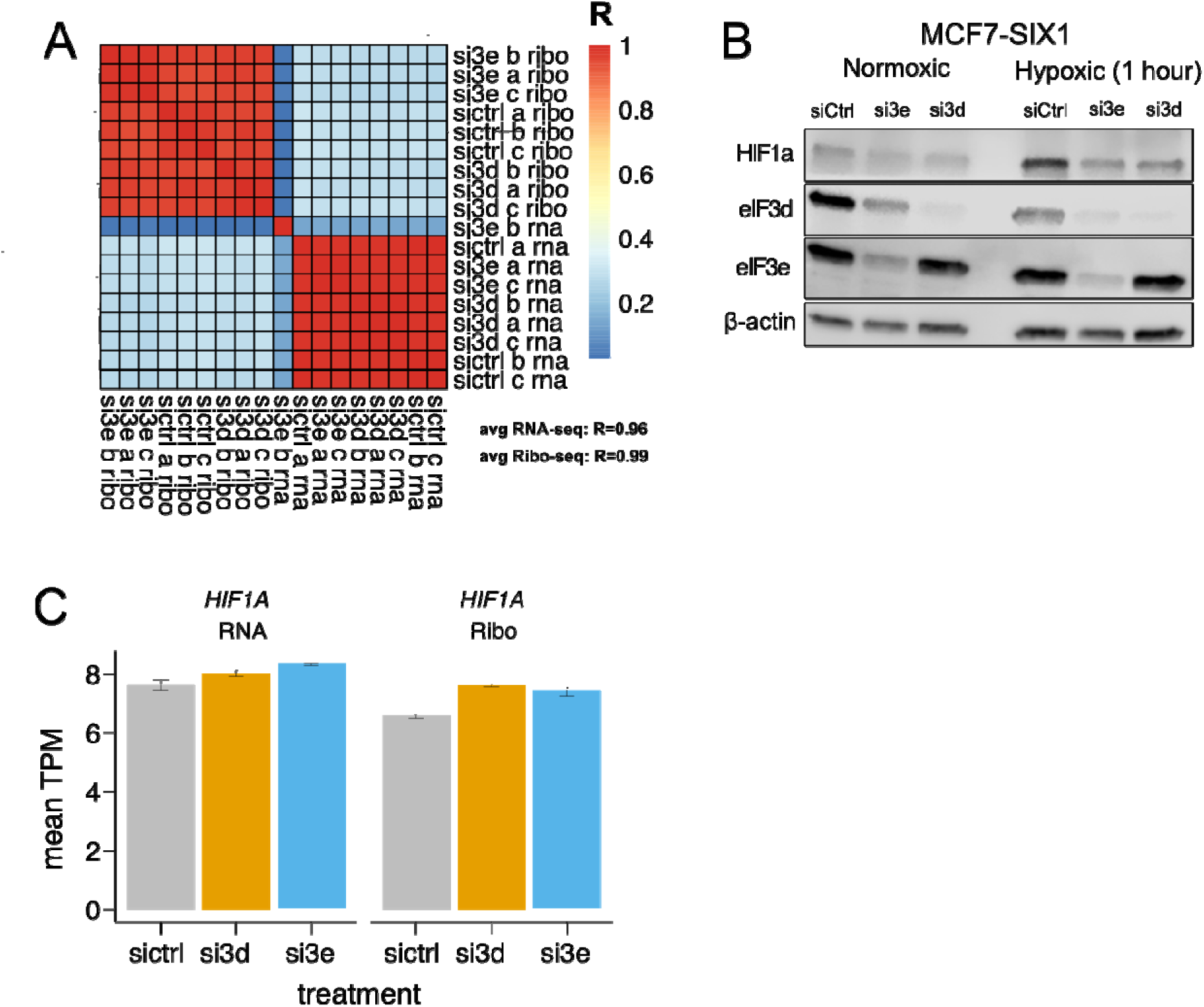
A. Heatmap of Pearson correlation coefficient for normoxic RNA-seq and Ribo-seq Iog2 transcript per million (TPM) of hypoxia-treated RNA-seq and RIBO-seq samples. Si3e rep b was identified as an outlier and removed from the analysis. B. Western blot analysis showing HIF1a after elF3d and elF3e knockdown in normoxic and hypoxic (1 hour at 1% O2) conditions. C. HIFIalpha RNA and ribosome footprint TPM per sample.

**Figure S3:**
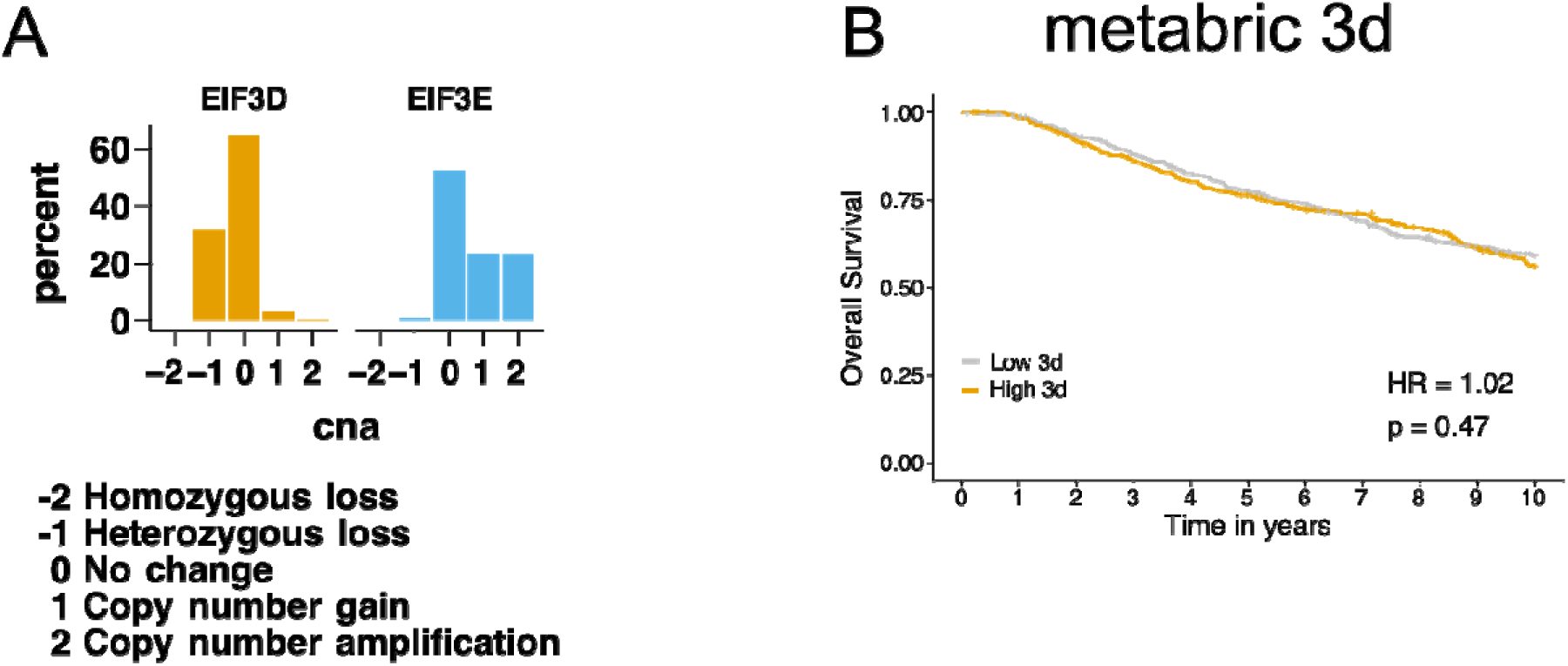
A. Copy number alterations (CNA) for EIF3D and EIF3E among METABRIC patients (n=1980). B. METABRIC patients stratified by enrichment/depletion of our elF3d signature. First (n=495) and fourth quartiles (n= 495) are visualized.

**Figure S4:**
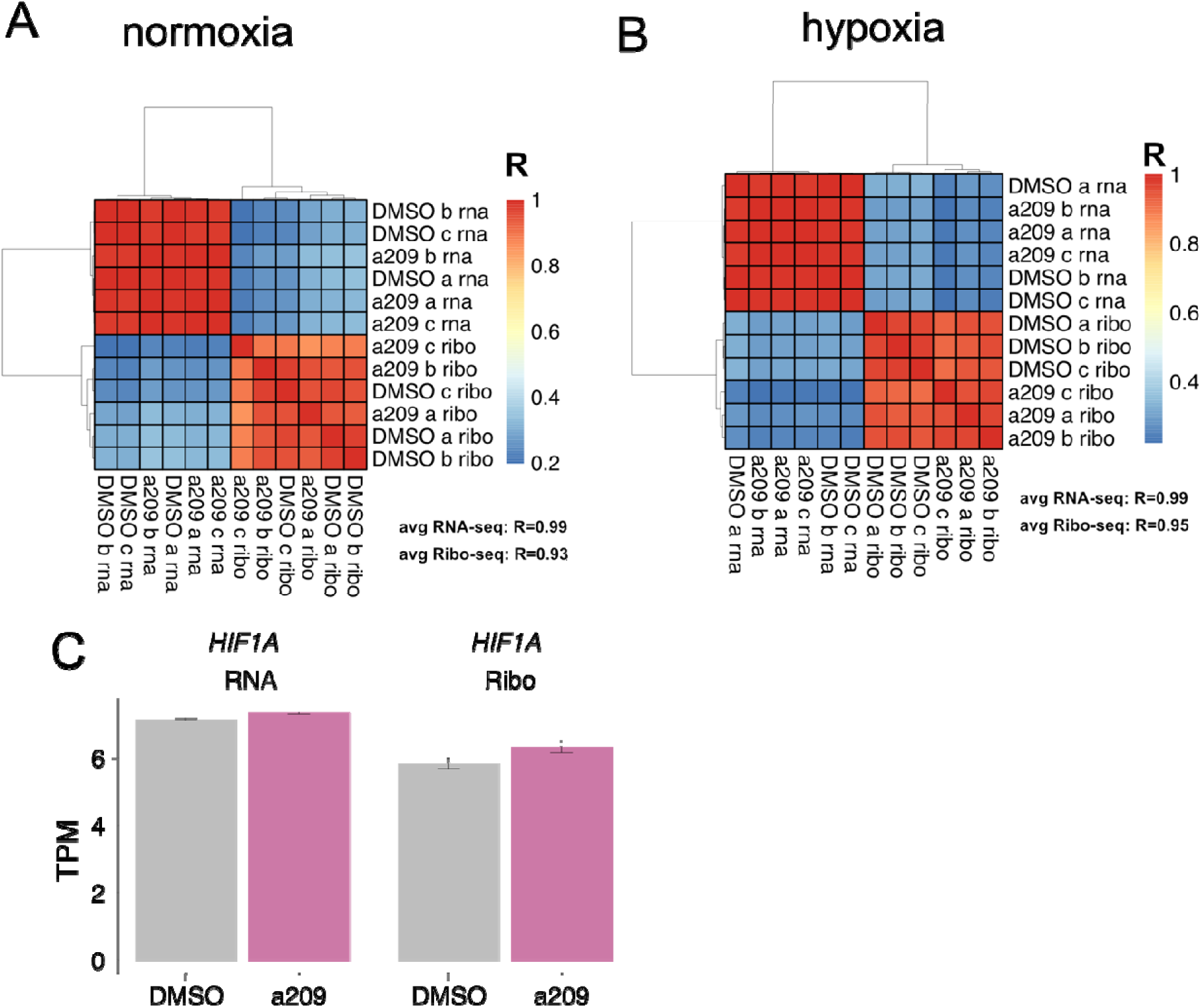
A. Heatmap of Pearson correlation coefficient for A) normoxlc and B) hypoxic RNA-seq and Ribo-seq Iog2 transcript per million (TPM) from DMSO or 209 treatment. 0) RNA and RPF TPMs of HIF1A with DMSO or 209.

**Figure S5:**
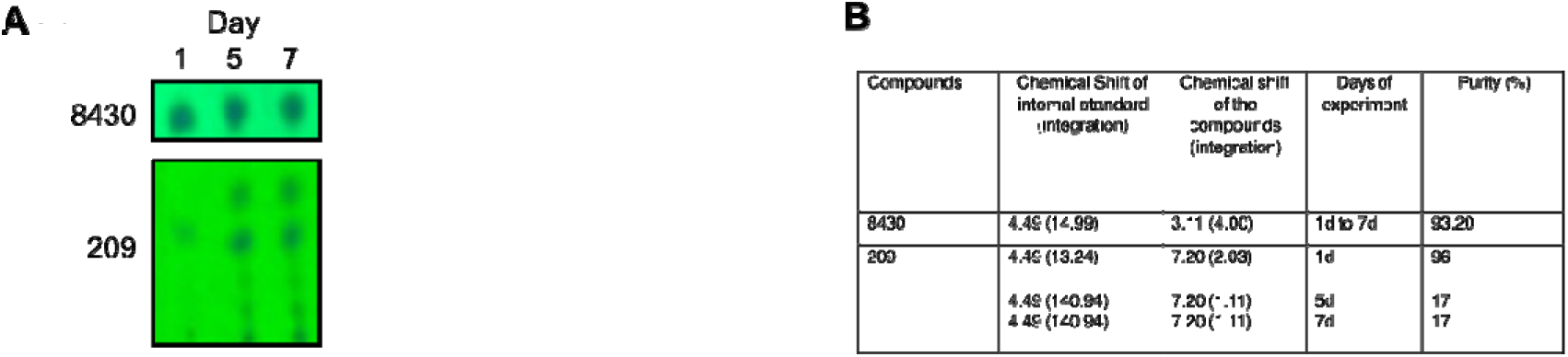
**A.** Thin layer chromatography (TLC) of 8430 after 1 day. 5 days, and 7 days. B. Quantitative analysis of 8430 and 209 in buffers with known quantity of ethelyn carbonate as internal standard for 1 d, 5d and 7days. Relative amount of compound at each day measured using qNMR.

**Figure S6:**
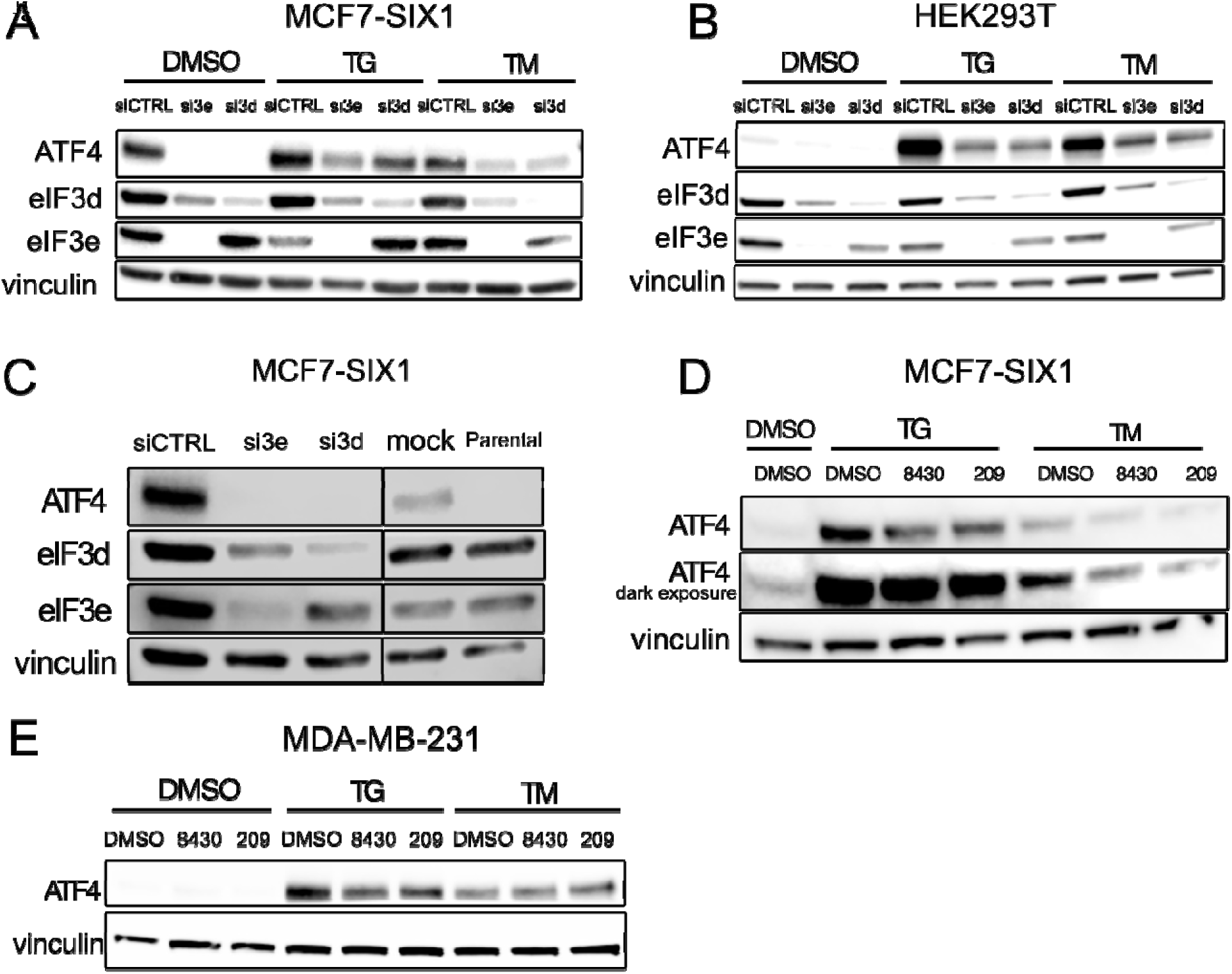
A. Western blot analysis showing ATF4 levels +/- TG treatment 100nM for 4 hours) or +/- TM treatment (2.5ug/mL for 4 hours) with elF3e or elF3d KD in MGF7-SIX1 cells and B. HEK293T cells. D. Western blot analysis showing ATF4 levels +/- TG treatment (100nM for 4 hours) or +/- TM treatment (2.5uM for 4 hours) with 8430 or 209 pre-treatment (20uM for 24 hours) in MCF7SIX1 cells or E. MDA-MB-231 cells.

**Data S1.** (separate file “Supplemental Datasets”)

| Sheet | Name | Figure | Guide |
| --- | --- | --- | --- |
| 2 | complete_te_list | calculate_te | Normalized TE, RNA, and RIBO counts for each gene in each condition |
| 3 | normoxia_sictrl_3e | 1 | Delta TE eIF3e KD compared to siCTRL normoxia |
| 4 | normoxia_sictrl_3d | 1 | Delta TE eIF3d KD compared to siCTRL normoxia |
| 5 | norm_h_table | 1 | Normoxia hallmark pathway enrichment for eIF3e KD compared to siCTRL and eIF3d KD compared to siCTRL |
| 6 | hypoxia_sictrl_3e | 2 | Delta TE eIF3e KD compared to siCTRL hypoxia |
| 7 | hypoxia_sictrl_3d | 2 | Delta TE eIF3d KD compared to siCTRL hypoxia |
| 8 | all_norm_to_hyp_changes | 2 | Delta TE normoxia to hypoxia for KDs and siCTRL |
| 9 | rna_siganture_3e_normoxia | 3 | RNA siganture siCTRL compared to eIF3e KD in normoxia (3e up) |
| 10 | rna_siganture_3d_normoxia | 3 | RNA siganture siCTRL compared to eIF3d KD in normoxia (3d up) |
| 11 | rna_siganture_3e_hypoxia | 3 | RNA siganture siCTRL compared to eIF3e KD in hypoxia (3e up) |
| 12 | rna_siganture_3d_hypoxia | 3 | RNA siganture siCTRL compared to eIF3d KD in hypoxia (3d up) |
| 13 | normoxia_dmsa_a209 | 5 | Delta TE a209 compared to DMSO normoxia |
| 14 | hypoxia_dmsa_a209 | 5 | Delta TE a209 compared to DMSO hypoxia |
| 15 | all_norm_to_hyp_changes_compound | 5 | Delta TE normoxia to hypoxia for a209 and DMSO |

## Notes

### Competing Interest Statement

The authors have declared no competing interest.

